# Annoying noise: effect of anthropogenic underwater sound on the movement and feeding performance in the red cherry shrimp, *Neocaridina davidi*

**DOI:** 10.1101/2022.10.10.511615

**Authors:** Sasan Azarm-Karnagh, Laura Lopez Greco, Saeed Shafiei Sabet

## Abstract

Acoustic pollution in aquatic environments has increased dramatically, with adverse effects on many organisms. Benthic organisms, including many invertebrates, can sense underwater sounds, yet the responses they trigger in these organisms have received little attention. This study investigates the impact of underwater sound on the behaviour of the red cherry shrimp *Neocaridina davidi* as a model of freshwater decapod. The effect of underwater sound exposure on the movement behaviour and feeding performance of individual shrimps was assessed. Movement speed decreased significantly upon opening the divider in both the sound and control treatments. However, there were no significant changes in total minutes between the control and sound treatments, implying no sound-related initial changes for releasing movement. The spatial distribution of shrimps in response to the sound treatment showed significant changes; shrimps spent more time at the farthest point from the sound source. The time to find the food source (latency) also increased in the sound treatment compared to the control. Moreover, in terms of the number of successes and failures in finding the food source in the control treatment, significantly more shrimps succeeded in finding the food source. Besides, the number of revisits to the food source decreased in sound treatment compared to control and more shrimps were significantly distracted in sound treatment. Our study highlights the crustacean’s ability to receive human-made sound. Thus, they are prone to the impacts of anthropogenic sound, causing negative impacts on their movement-swimming activities, and feeding behaviour and exposing them to potential predator threats. Affecting foraging performance in this gregarious species may have detrimental impacts on their reproductive success and, subsequently unexpected fitness consequences.

## Introduction

Over recent decades, environmental pollution caused by human activities has altered terrestrial and aquatic habitats globally (Kight and Swaddle, 2011). Aquatic animals use signals and exploit cues to explore their habitat and perform various biological activities (Kight and Swaddle, 2011). In aquatic habitats, anthropogenic sensory pollutants such as chemical contaminants (Tornero and Hanke, 2016), artificial light at night (ALAN) (Gaston et al., 2015) and anthropogenic sound (Kunc et al., 2015, Peng et al., 2015) have quite diversely changed the chemical components, daylight luminance and acoustic characteristics of the environment globally, and subsequently encountered many species under threat of extinction. All these sources of pollutants are prominent and pervasive and are perceived by aquatic animals via different sensory mechanisms, modalities, and organs. Animals rely on their sensory organs in different modalities (e.g. smell, vision and hearing) to perceive and process a range of information (signals and cues) in their habitats (Ukaogo et al., 2020). The ability to exploit such habitat information is crucial and a prerequisite for survival. Interference with these sensory systems due to introducing novel stimuli may be detrimental to their feeding efficiency success, communication with con-specifics, locating prey, avoiding predators, and courtship performances (Dominoni et al., 2020). Sound, as an informative sensory stimulus, is a key signal in the aquatic environment because it is not attenuated as fast as chemical or light pollutants and can be propagated over long distances (Hawkins and Myrberg, 1983).

Sound pollution is omnipresent in aquatic habitats and may affect large areas, therefore negatively affecting individual species, as well as changing species interactions and animal communities (Kunc et al., 2016; Shannon et al., 2016; Slabbekoorn et al., 2010, 2019, De Jong et al., 2018). Although the underwater environment has long been considered a “silent world” (Cousteau and Dumas, 1987), recent studies have revealed that it is full of sounds and vibrations (Williams et al., 2015). These can arise from natural sources such as waves, rain and currents or from biotic sources such as animal vocalization, displacement, hunting and feeding (Gordon and Tyack, 2001; Stocker, 2002; Scott, 2004). Anthropogenic activities also contribute to underwater sound and vibrationincluding construction, resource extraction, transportation, aquaculture, fishing and recreation (Slabbekoorn et al., 2010). The marked increase in anthropogenic sound and associated vibration has changed the underwater acoustic landscape (McDonald et al., 2006; Andrew et al., 2011; Frisk, 2012; Duarte et al., 2021). Shipping noise, motorboat sound, and recreational boating are the main sources of anthropogenic sound in most of the world’s oceans (Hildebrand, 2009) and freshwater habitats (Mickle & Higgs, 2018), with a typical continuous acoustic temporal pattern being the most dominant within the low-frequency portion of the spectrum (propulsion systems of ships: <200 Hz). Moreover, pumping systems, aerators and feeding machines produce continuous sound patterns in fish facilities, laboratories, and ornamental pet shop centres (Lara & Vasconcelos, 2019; Radford & Slater, 2019).

Research on the impact of anthropogenic sound in the aquatic environment has focused on marine mammals (Androw et al., 2007;; Nowacek et al., 2007; Southall et al., 2008) and fish (Bruintjes et al., 2015; Simpson et al., 2016; Herbert-Read et al., 2017), while the potential impacts on invertebrates have received less attention (Wale et al., 2013; Roberts et al., 2015, 2016; Williams et al., 2015). Although invertebrates are quite diverse, important for all trophic levels, and have a relatively large abundance in marine and freshwater environments (Morley et al., 2014; Solan et al., 2016), their capability to detect and potentially exploit sound is relatively unknown. Marine decapods play an important role in top-down control of prey populations and can shift the abundance and community structure in aquatic habitats (Reise, 1979, Bell and Coull, 1978). Thepotential impacts of sound may cause cascading impacts on many species in a variety of taxa (Slabbekoorn et al., 2010; Shafiei Sabet et al., 2016a,b). In contrast to marine habitats, given the broad-ranging diet of the red cherry shrimp, it is likely that this species also influences food webs in freshwater habitats. The potential effects of anthropogenic sound on freshwater decapods and their role in regulating top-down and button-top control of predator-prey interaction are unknown.

Elevated sound levels could affect foraging behaviour in three ways, which are not mutually exclusive. First, noise could act as a stressor (Williams et al., 2015), decreasing feeding behaviour directly through reduced appetite (Charmandari et al., 2004), or indirectly through a reduction in activity and locomotion (Mendl, 1999). It can also alter the cognitive processes involved in food detection, classification and decision-making (De Kloet et al., 1999; Lupien & McEwen, 1997). Second, noise could act as a distracting stimulus, diverting an individual’s limited attention from their primary tasks to the noise stimuli now present in the environment (Chan & Blumstein, 2011; Mendl, 1999). This could impair foraging success if, for instance, suitable food sources are detected less often or more slowly, are assessed less accurately, or if prey items are mishandled (Purser & Radford, 2011). Third, noise could mask crucial acoustic cues (Brumm & Slabbekoorn, 2005). If cues produced by prey are masked, feeding opportunities may be missed (Schaub et al., 2008; Siemers & Schaub, 2011). If acoustic predator cues are masked and animals compensate by relying on visual information (Quinn et al., 2006), then visually guided food searching and acquisition may be compromised.

The red cherry shrimp (*Neocaridina davidi*) is a tiny and gregarious benthic freshwater invertebrate species of shrimp with a wide distribution, occurring from Southeast Asia to Europe (Wowor et al., 2004; Liang, 2004; Klotz et al., 2013, Onuki & Fuke, 2022). Although this species is an ornamental crustacean and provides an important link between trophic levels and food webs (Kelly et al., 2012, Weber and Traunspurger, 2016), its behavioural responses to stress stimuli are still poorly understood.

In this study, two experiments were conducted to analyse the effect of sound exposure on the locomotor behaviour and feeding activities of the red cherry shrimp under laboratory conditions. The first experiment evaluated the initial releasing movement speed, spatial distribution, food-finding latency (the time to find the food source), the number of successes and failures in finding the food source, and the number of revisits to the food source in response to ambient sound conditions (control treatment) and an elevated generated sound level. The second experiment investigated sound-related feeding distraction in response to ambient sound conditions as a control treatment and an elevated generated sound level. Red cherry shrimps were exposed to a 400-2000 Hz acoustic stimulus mimicking a frequency bandwidth range that they can detect and which represents the frequencies produced by human activities that shrimps are exposed to in their natural habitats and at rearing facilities (Lovell et al., 2005; Slater et al., 2020).

## Material and methods

### Ethical Note

We adhered to the Guidelines for the treatment of animals in behavioural research and teaching (ASAB, 2020). There are no legal requirements for studies involving crustaceans in Iran.

### Study Animals and Husbandry

All shrimps were purchased from a local aquatic pet dealer in Rasht city, Iran. For acclimation to the new tank conditions, the animals were transferred to a glass holding tank (Figure 1) (48 × 28 × 32 cm and 4 mm wall thickness) for one week equipped with a 150 watts heater, sponge biofilter, Java moss, natural plant, a 12 watts LED light, and sandy bed and plant soil. Water ° temperature was kept at 26±1 °C, aeration 5L/min, and the light period was 14 D:10 L. Water quality was maintained within safe parameters. Shrimps were fed every 12 hours with *Spirulina* algae tablets, with any excess cleared from the holding tank during tank maintenance no more than 4 h after feeding. Although there was a constant water change within the holding tank, 25% of water was removed by siphon with excess food and waste; this water was replaced by normal tank flow-through. Water changes and the flow-through system ensured the maintenance of constant oxygen levels and the removal of any metabolic waste materials. The physical and chemical conditions, such as ambient acoustics, water temperature, water changes, and lack of stress hormones for both tanks (holding and experimental), were similar. The sex ratio of shrimp was 1:1, and all mature shrimps had a body length ranging from 2 to2.5cm.

**Fig. 1.**
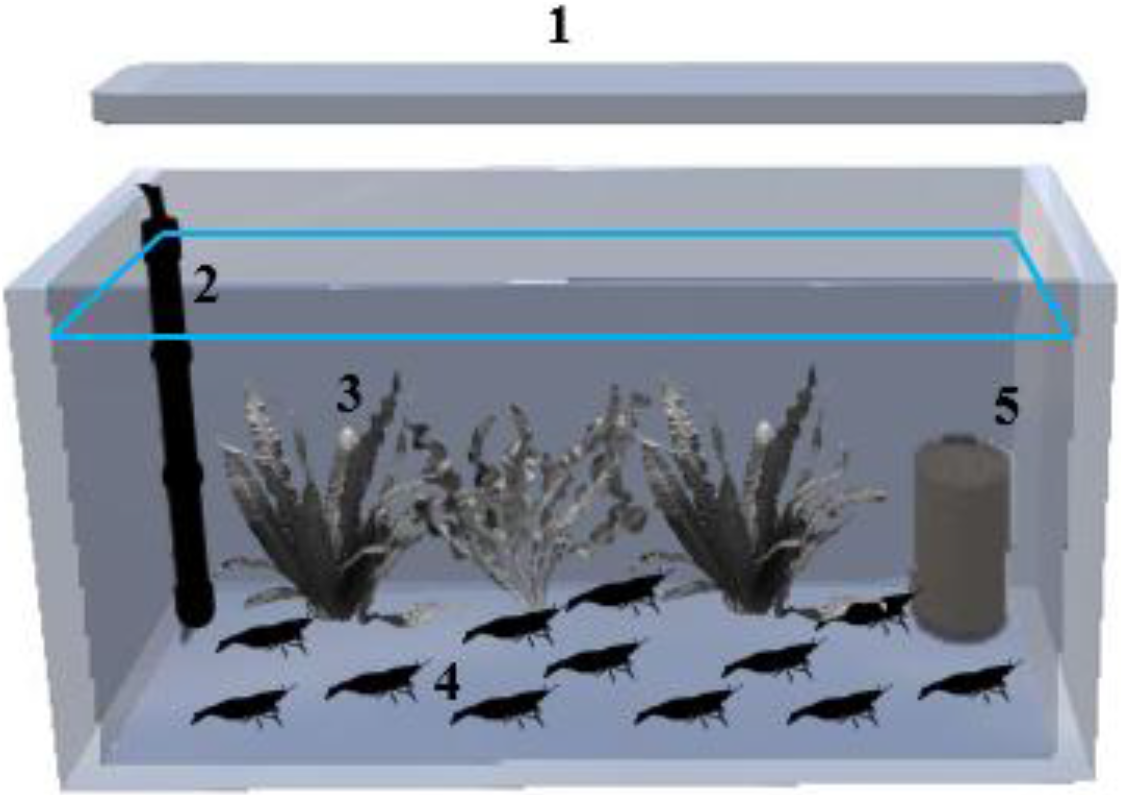
Experimental configuration to maintain *Neocaridina davidi* in the holding aquarium. Set up of the holding tank: 48 × 28 × 32 cm and 4 mm wall thickness. 1) LED; 2) heater; 3) aquatic plant; 4) shrimps, and 5) sponge filter.

### General Experimental Design

Both experiments were conducted in a tank (40 × 30 × 30 cm) with 4 mm thick walls, two divider plates, and a dewatering surface of 20 cm. Water temperature, pH, total dissolved solids (TDS), light intensity, and NO_3_^-^, NO_2_^-^, NH_4_^-^were measured (Table 1).

**Table 1.** The physic-chemical measurement of water

| Parameters/value | Amount | Device/model |
| --- | --- | --- |
| Temperature ($^{\circ}\text{C}$) | $26 \pm 1$ | Mercury thermometer |
| Light intensity (Lux) | $113.8 \pm 2$ | Lux meter (model: Digital Lux Meter-GM1010) |
| TDS (ppm) | $160 \pm 1$ | TDS meter (model: TDS-3) |
| pH | $7.2 \pm 2$ | pH meter (model: SANA SL-901) |
| $\text{NO}_3^-$ (mg/l) | $<0.2$ | Bante321-$\text{NO}_3^-$ Portable Nitrate Ion Meter |
| $\text{NO}_2^-$ (mg/l) | 0 | JBL Nitrite Test Kit |
| $\text{NH}_4^+$ (mg/l) | 0.21 | Bante321-$\text{NH}_4^+$ Portable Ammonium Ion Meter |

Light intensity was recorded daily at the beginning and end of each treatment. Light intensity was measured 20 cm above the water level and in the bottom-middle of the tank. The other parameters were measured daily before each treatment. In each experiment, 35 shrimps were used for each treatment with an alternation of both males and females. To eliminate the effects of test time on the treatments, the order of control and sound treatments were applied one by one. The playback speaker was placed behind a vertical partition 6 cm from one short side of the tank. The experimental tank was placed on a 5 cm thick foam pipe lagging to minimize sound transfer and it’s sides were covered with opaque black paint to minimize distraction from visual reflections.

### Sound characteristics

The sounds used in this study were created using specialized software (Audacity-win-vr 2.3.1), based on Shafiei Sabet et al. (2015). The ambient sound frequency (control) was 400 to 2000 Hz with anintensity of 96.54±1 dB in WAV format. For the elevated sound treatment, a sound with a continuous pattern and frequency of 400 to 2000 Hz, intensity of 110.40 ±1, and WAV format was used. The continuous sound with this frequency range was chosen due to its’ similarity to the sound of motor boats, aeration pumps, and filtration systems that are commonly used on farms (Peng et al., 2015). A 30-watt hand-built underwater speaker with a frequency of 0 to 3000 Hz, a model laptop (Lenovo-V110), an amplifier, and a 12 V sealed battery were used to supply the required power.

### Sound measurement and analysis

The sound emitted in the experimental tank environment was measured using a sound measuring device (TASCAM-DR-100MK2) equipped with a calibrated hydrophone by a specialized standard method (Shafiei Sabet et al., 2015). The sound data were imported into R (v.3.6.3) for analysis under specialized acoustic programming and the sound intensity (dB ref 1μPa), frequency (Hz), sensitivity and sample rate were measured. Graphs of sound spectrum that indicates the amount of sound energy (Figure 2) and sound pressure level or SPL per unit (dB ref 1μPa2 / Hz) were drawn by the software (Figure 3).

**Fig. 2.**
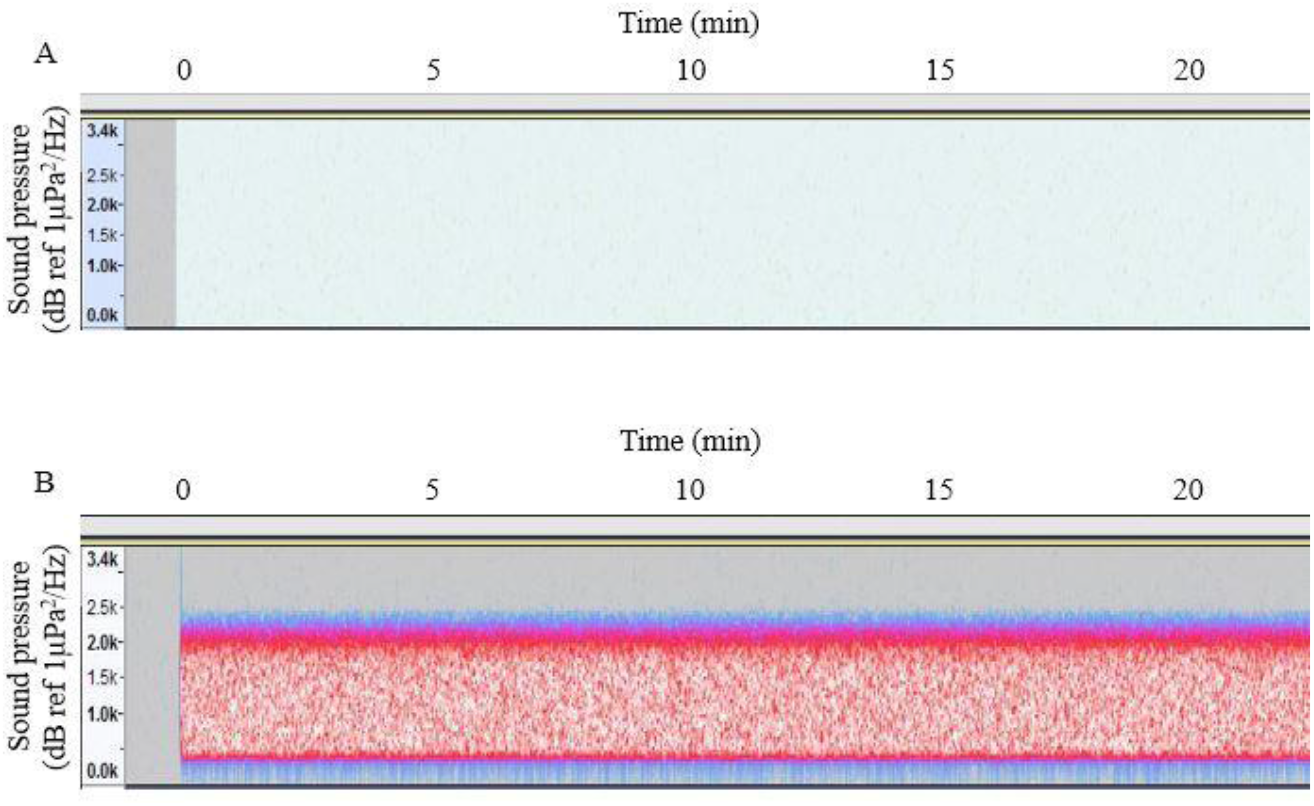
Spectrogram (for 20 s time slot) magnification of A) Control and B) Sound wave (Sound treatments) used in the experiment: frequency (kHz) vs. time (min). The intensity is reflected by the colour scale (dB re 1 lPa ^2^/Hz).

**Fig. 3.**
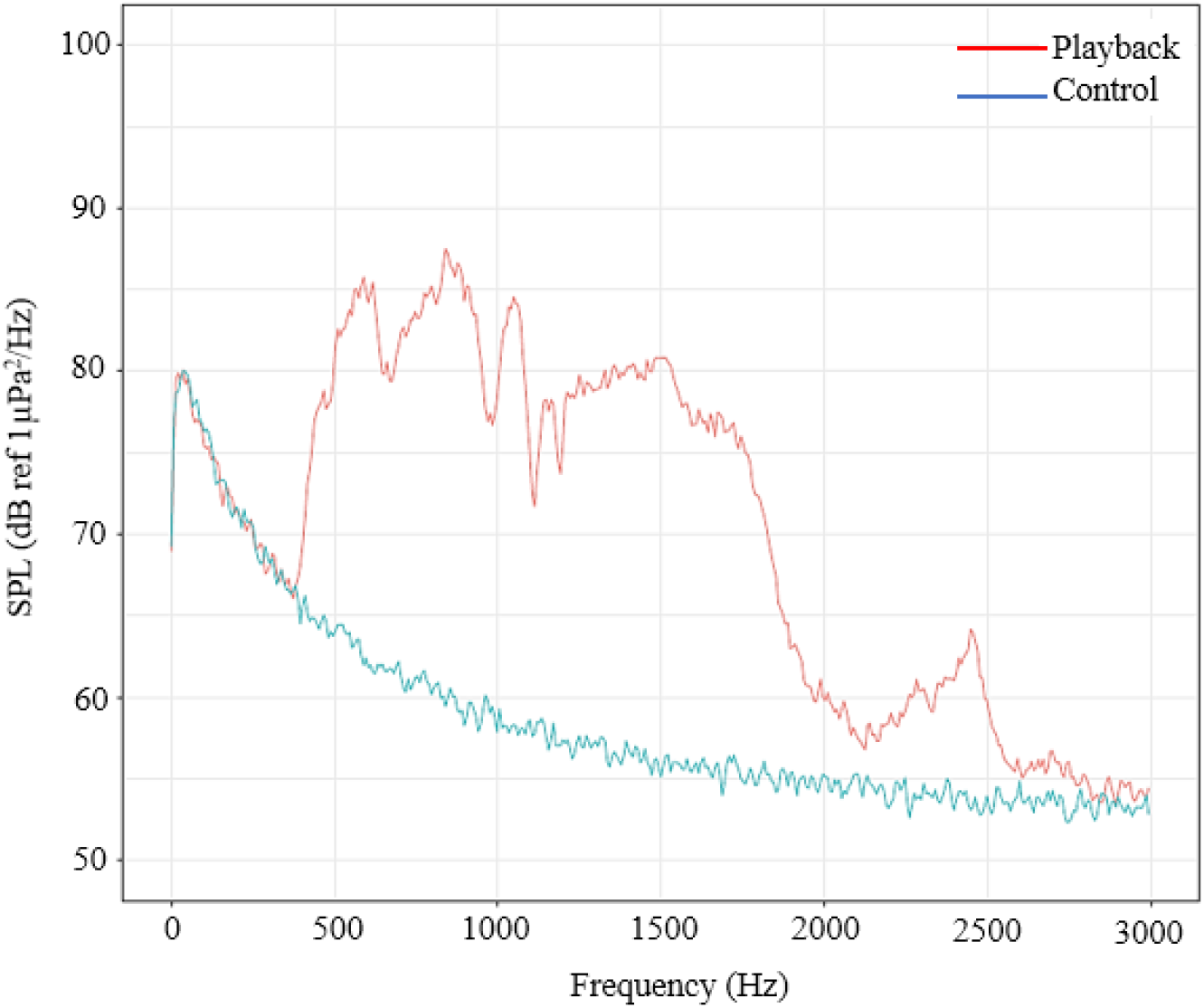
Mean power spectrum of the sound stimuli (red line) and of the experimental tank background noise (blue line). Sound Pressure Level expressed in dB re 1 μPa^2^/Hz (Welch’s method, Hann window, FFT length 1024).

### The first experiment: movement speed, spatial distribution, food-finding latency, number of revisiting and food finding

In this experiment, before starting the main experiments, shrimps (n = 70; 35 individuals per treatment) were denied access to food for 48 hours in the holding tank. Then, in order to acclimation to the experiment tank (Figure 4), 1 hour before the test, they were individually transferred to the starting area behind a divider plate. All transferring procedures were gently and same for trials. At this time, the food source was added to the other side of the divider plate (see Figure 4). In the control treatment, the ambient noise was played (20 minute duration)at the same time as the divider plate was removed (2 cm upwards from the bottom of the aquarium, with minimal physical movement of the divider plate so that less stress is applied to the shrimp and it can pass through it) (Figure 5a). Each shrimp was given a maximum of 10 minutes to find the food source. The sound treatment (20 minute duration) played at the same time as the divider plate was removed and each shrimp was given a maximum of 10 minutes to find the food source (Figure 5b). The behavioural indicators measured are described in Table 2.

**Fig. 4.**
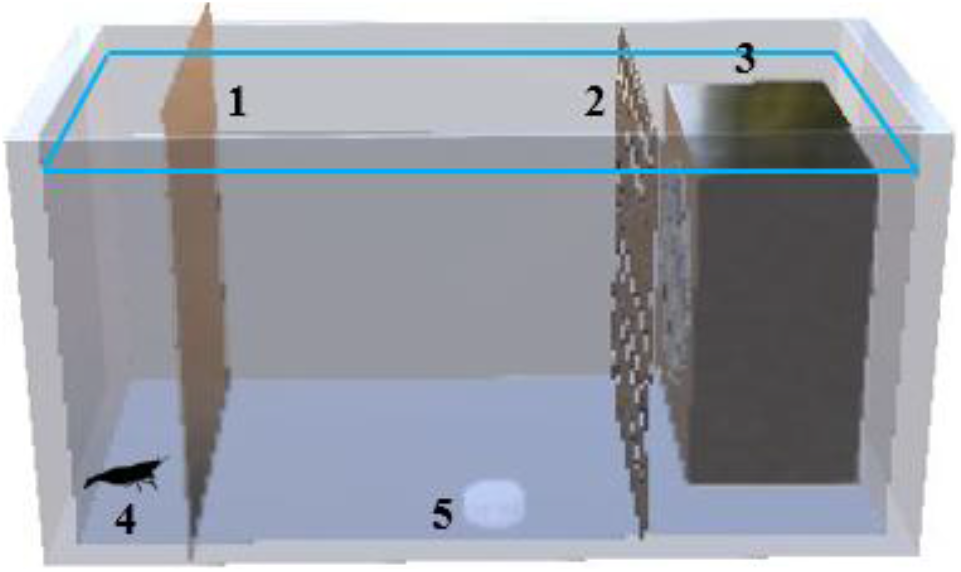
First experiment set up (40 × 30 × 30 cm), 1) divider, 2) speaker holder, 3) underwater speaker, 4) the shrimp, 5) food source.

**Fig. 5.**
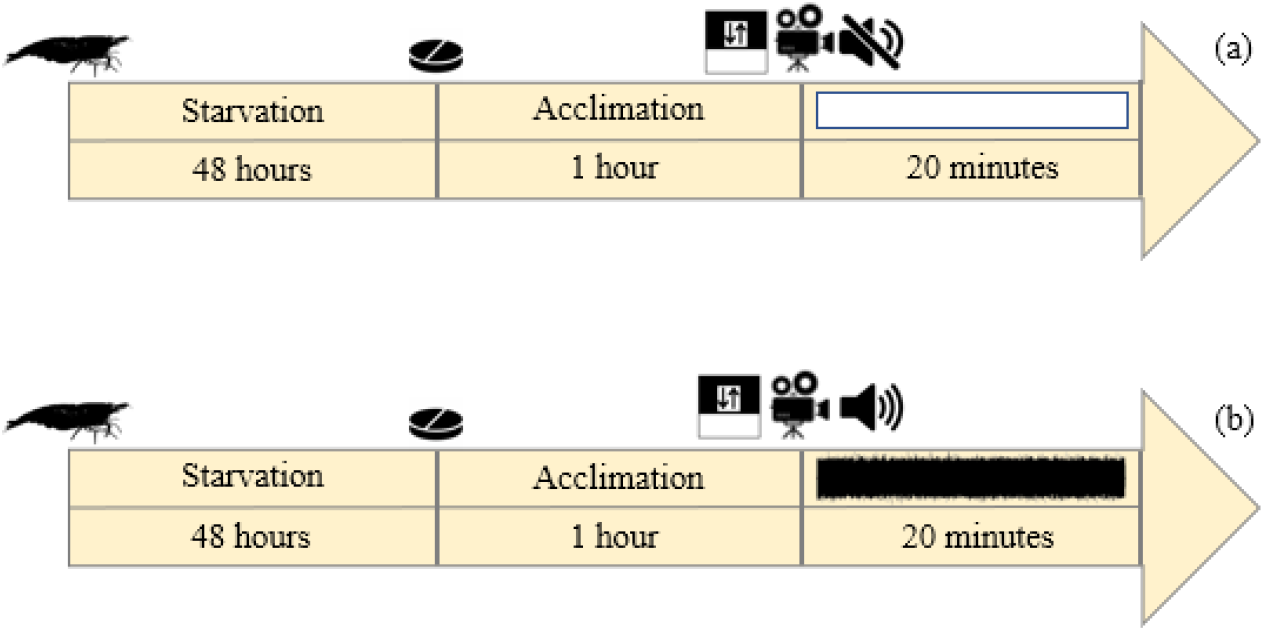
Timeline view of the first experiment in a) control and b) sound treatments with *Neocaridina davidi*. 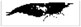 the shrimp,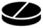 food source,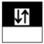 opening divider, 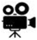 start filming,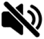 ambient noise,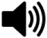 sound.

**Table 2.** Description of behavioural actions by the Red cherry shrimp (*Neocaridina davidi*) registered during the experimental tests, based on videotapes visualizations.

| Behaviour | Description | Unit | Application test |
| --- | --- | --- | --- |
| Movement speed | Mean distance moved by the center point of the subject per unit time | Centimetres/seconds | Single |
| Spatial distribution | The mean attendance of individuals in a specific area | Seconds & Centimetres | Single |
| Food-finding | Number of individuals that succeed to find the food source | Number of individuals | Single |
| Feeding latency | Mean time to find the food source | Seconds | Single |
| Number of revisiting | Number of revisiting to food source for a certain period of time | Number of individuals | Single |
| Feeding distraction | Number of individuals were complete freezing and movement away from the food after sound playback | Number of individuals | Single |

### The second experiment: feeding distraction

In this experiment, shrimps (n = 70; 35 individuals per treatment) were denied access to food for 48 hours in the holding tank. For both experiments, the size and sex ratio, feeding regime and feed restriction period were the same to avoid any confounding factors affecting behaviour unintentionally. In the control treatment, after 1 hour acclimation behind the divider plate in the experiment tank (Figure 6), the divider plate was removed and the shrimp were allowed to search for food without playing any sound. As soon as the shrimp found the food and started feeding, the control treatment (ambient noise) was played for 20 minutes and each reaction was recorded in real time by the observer (Figure 7a). In the sound treatment, after 1 hour of acclimation behind the divider plate in the experimental aquarium, the divider plate was removed and the shrimp were allowed to search for food without any sound (Figure 7b). As soon as the shrimp found the food and started feeding, the sound treatment was played for 20 minutes and each reaction was recorded in real time by the observer and by camera. A feeding distraction is defined as a shrimp stopping eating, freezing and moving away from the food source (within 10 minutes). Therefore, in this experiment, the number of shrimps whose feeding function was disturbed versus undisturbed was recorded in each of the control and sound treatments.

**Fig. 6.**
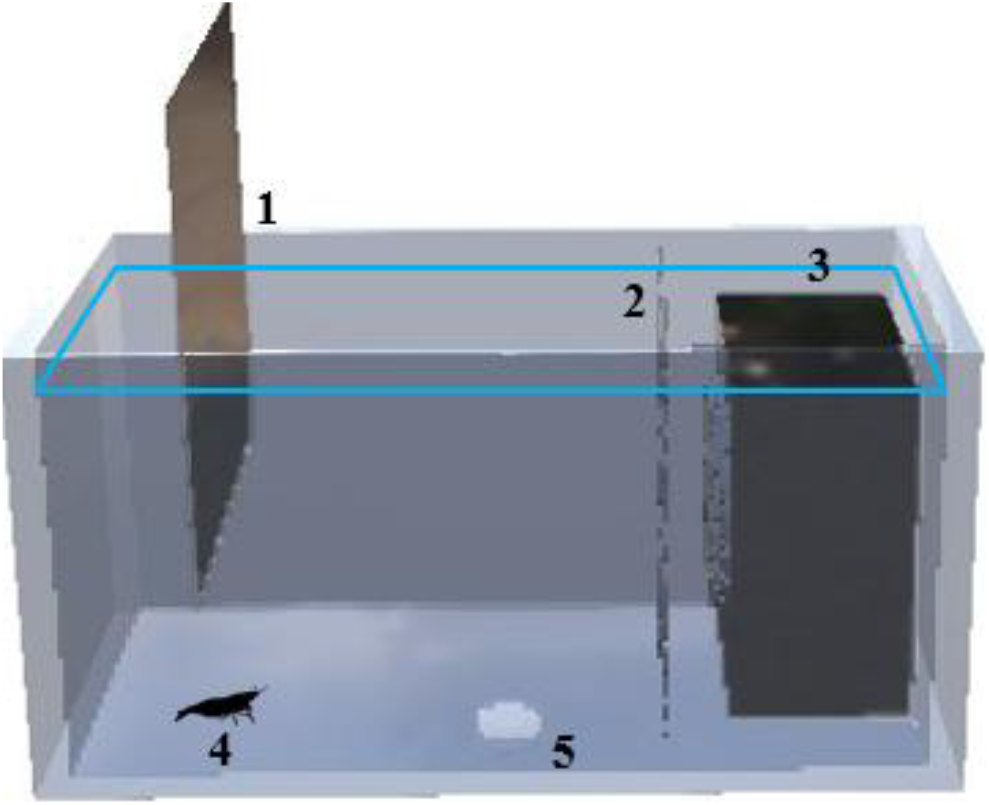
Second experiment set up (40 × 30 × 30 cm). 1) divider, 2) speaker holder, 3) underwater speaker, 4) the shrimp, 5) food source.

**Fig. 7.**
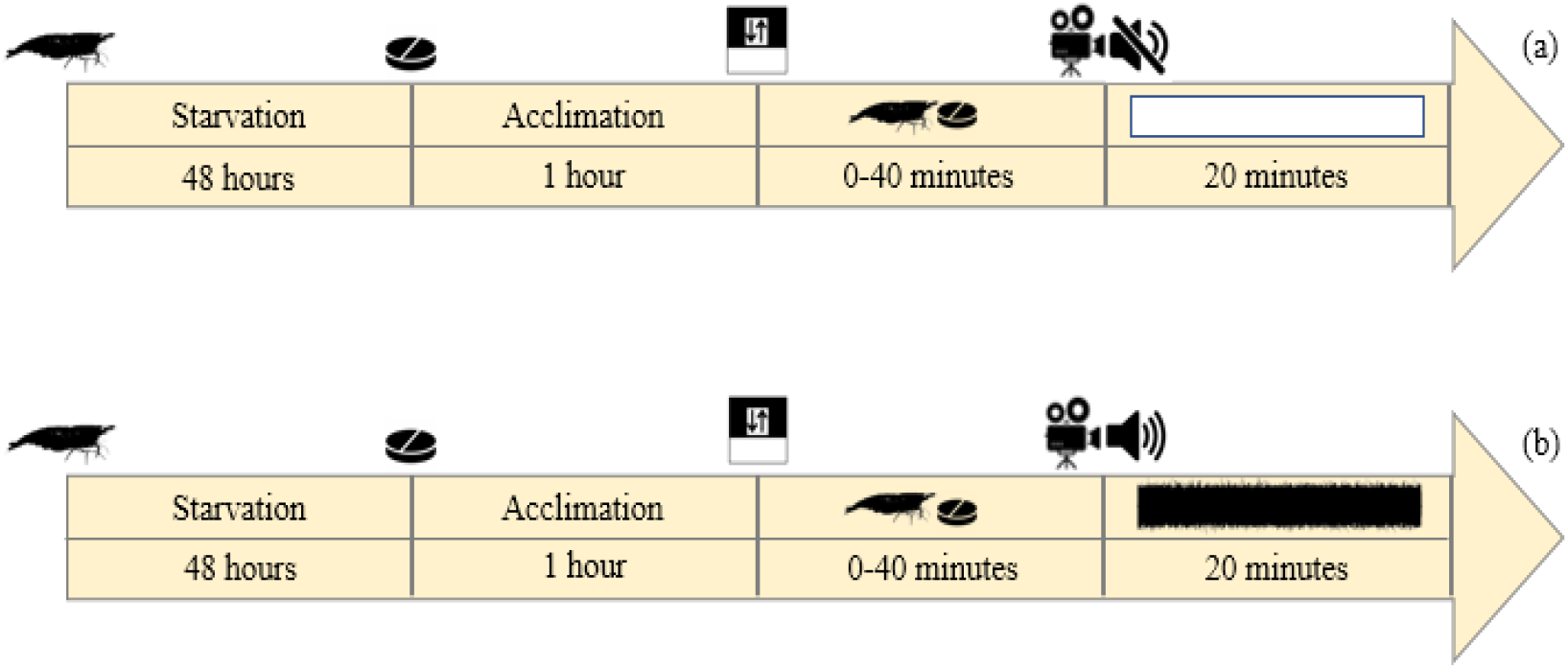
Timeline view of second experiment in a) control and b) sound treatments.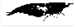 the shrimp,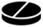 food source, 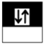 opening divider, 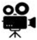 start filming,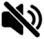 ambient noise,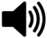 sound.

### Statistical Analysis

Statistical analysis of data was performed using SPSS-25 and Excel-2019 software. Kolmogorov-Smirnov test and Levene test were used to test the normality of the data. For movement speed, a one-way t-test and one-way repeated measures were used. For spatial distribution, revisiting and latency, an independent t-test was used at the level of 0.05. Chi-square test was also used for food-finding and feeding distraction. To analyze behavioural observations in all trials, the recorded videos were reduced to 1 fps magnification (due to the slow movement of the shrimp and the ease of recording the movement points) and the movement speed and spatial distribution of the shrimps were analyzed using Logger Pro 3.14.1 software.

## Results

### Movement speed

There were no significant changes between total movement speed for the control and sound treatments (Figure 8) (P>0.05). However, as soon as the divider was opened, the initial movement speed in the sound and control treatments, showed a significant decrease (P<0.001) (Figure 9).

**Fig. 8.**
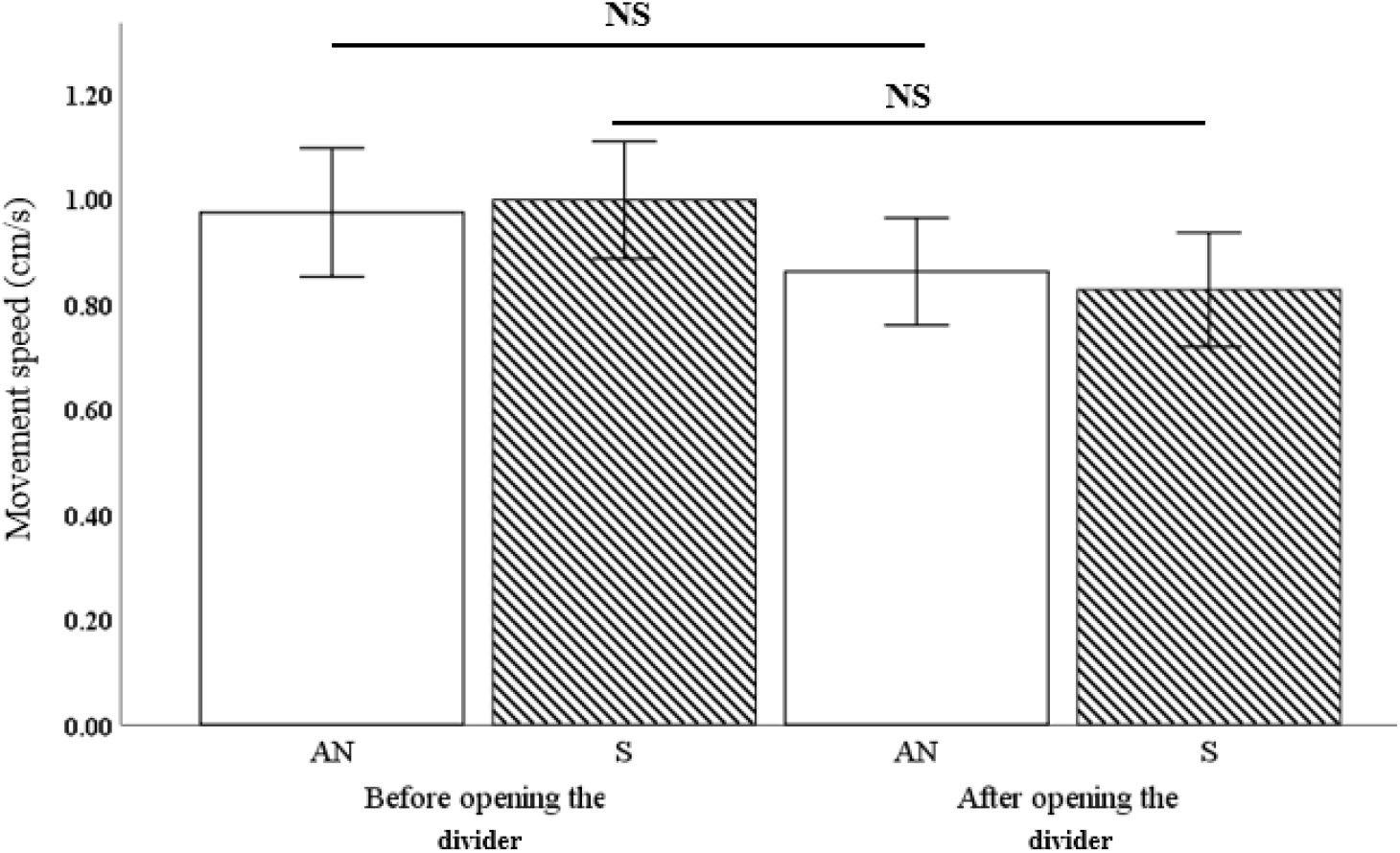
Mean ± SE movement speed (cm/s) for each sound treatment; ambient sound condition as control (AN) and sound treatment (S) for 35 red cherry shrimps. Red cherry shrimp movement speed did not change with sound conditions among treatments. NS: Not Significant.

**Fig. 9.**
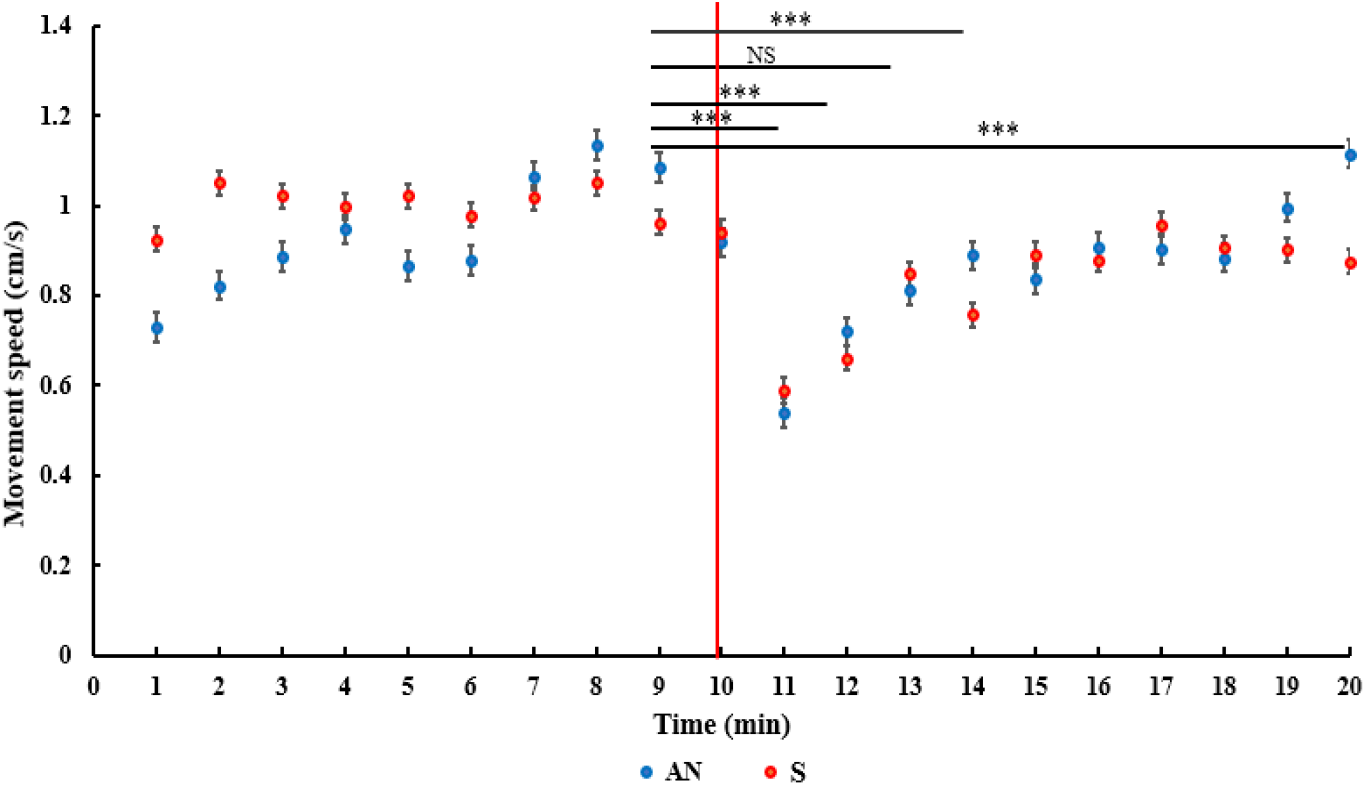
Movement speed in control (AN) and sound (S) treatments changed as soon as opening the divider. As soon as opening the divider significantly movement speed reduced. NS: Not Significant, ***P<0.001.

### Spatial distribution

Shrimps in the control treatment were equally distributed in the available space before opening the divider. After opening the divider and allowing the shrimp to enter the new environment they were again equally distributed throughout the whole space (Figure 10, A and B). In the sound treatment, before opening the divider, the shrimps were equally distributed in the available space, but when the divider was opened and the shrimp were exposed to the sound, they dispersed significantly to the left of the tank (farthest point from the sound source) (Figure 10, C and D).

**Fig. 10.**
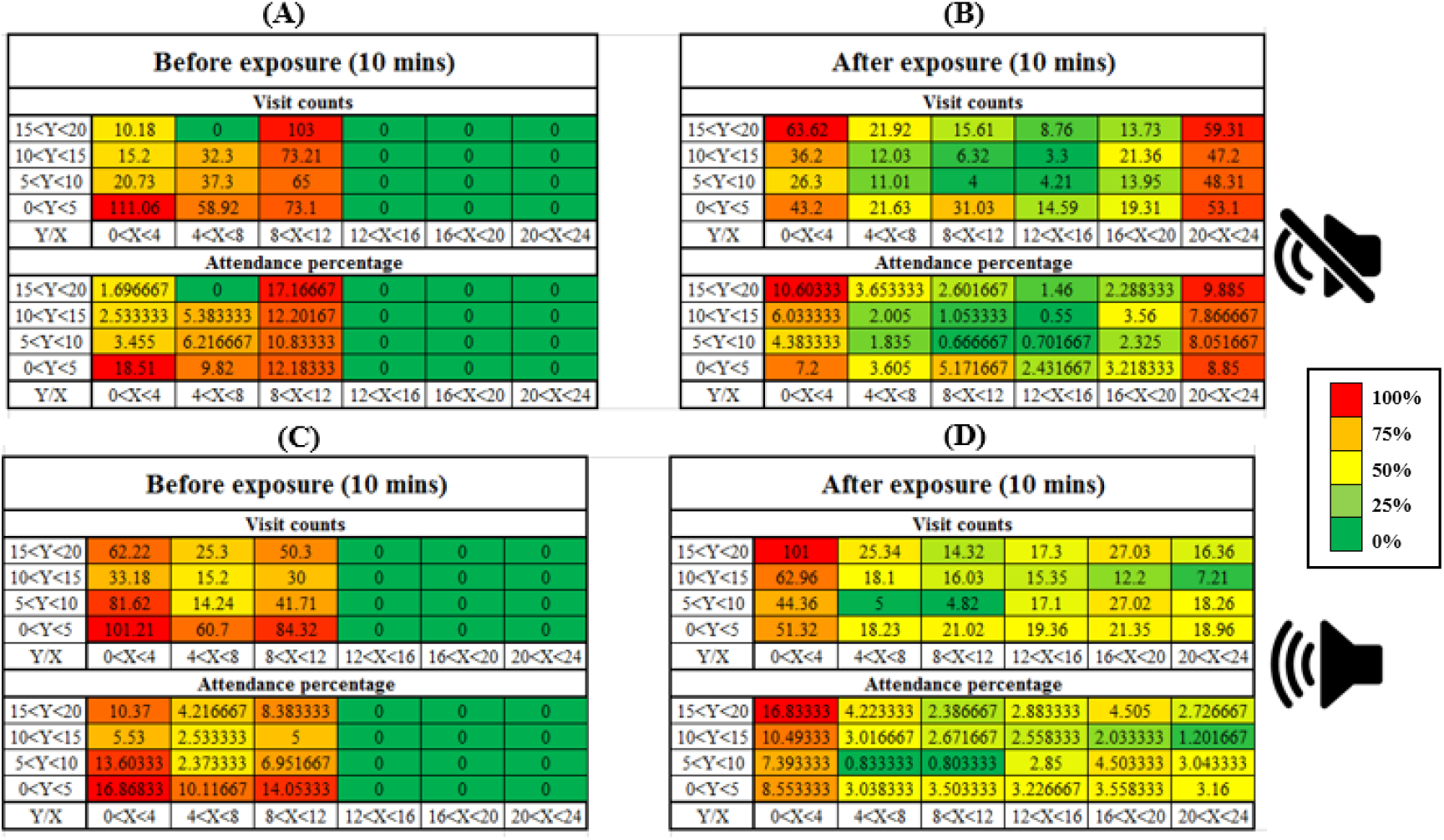
Cluster heatmap showing the spatial occupancy in terms of time spent in movement (expressed in seconds) by the shrimps among the aquarium cells. On the right wall, the location of the underwater sound speaker is presented. A,B) control condition C,D) sound condition.

Figure 11(A and B), shows the spatial distribution time of 70 shrimps in the right third of the tank (near the sound source) and the left third of the tank (away from the sound source). In the control treatment, the distribution of shrimp on both sides of the tank was the same and no significant changes were observed (P >0.05). In the sound treatment, the horizontal distribution of shrimp on the left side of the tank (the farthest distance from the sound source) significantly increased (P <0.01).

**Fig. 11.**
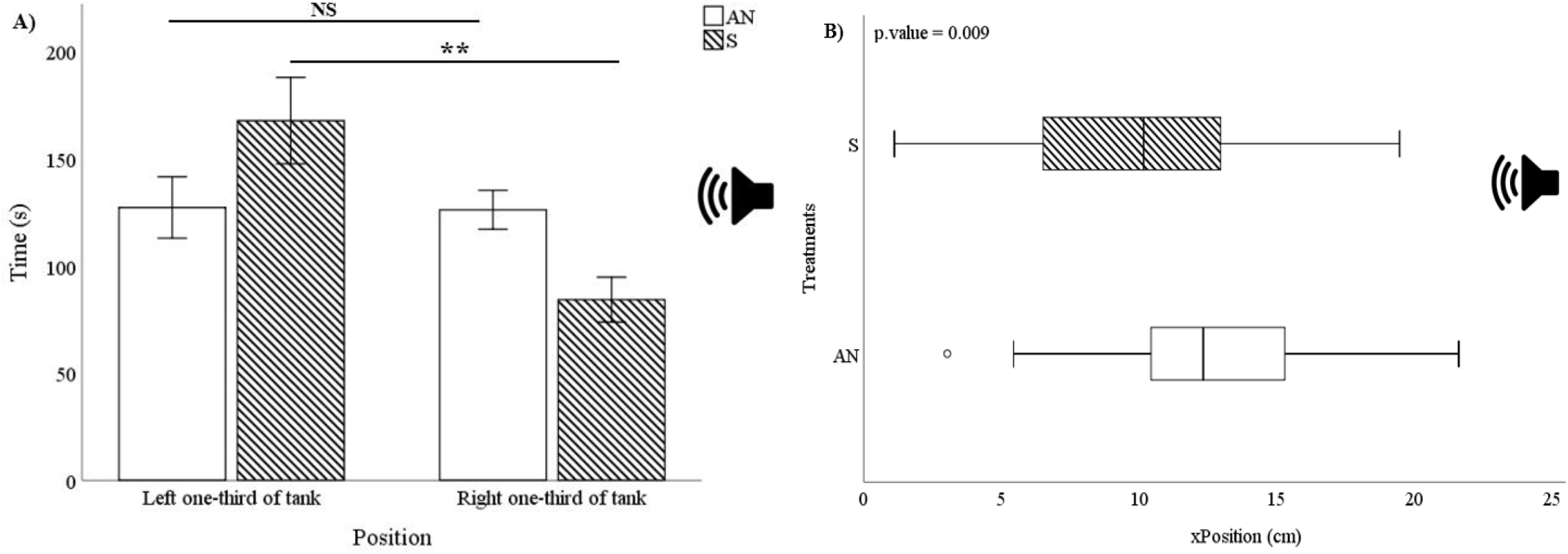
A) Mean ± SE time-place budget distribution of *Neocaridina davidi*. In sound condition the distribution of shrimp significantly increased in the left one-third of the tank (the point farthest from the speaker). B) Mean ± SE spatial distribution in horizontal (x) position in AN: Ambient Noise and S: Sound conditions, 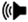 position of underwater speaker in tank. NS: Not Significant, **P<0.01

### Feeding activities

In the sound treatment, individuals found the food source significantly later than the control treatment (Figure 12 A, N=35, P <0.01). One individual in the control treatment and 9 in the sound treatment were not successful in finding the food source during the entire 10 minutes. The number of revisits to the food source was significantly reduced in the sound treatment (Figure 12 B, N=35, P <0.001), and significantly more individuals succeeded in finding the food source in the control treatment compared to the sound treatment (Figure 12 C, N=35, P <0.001)

**Fig. 12.**
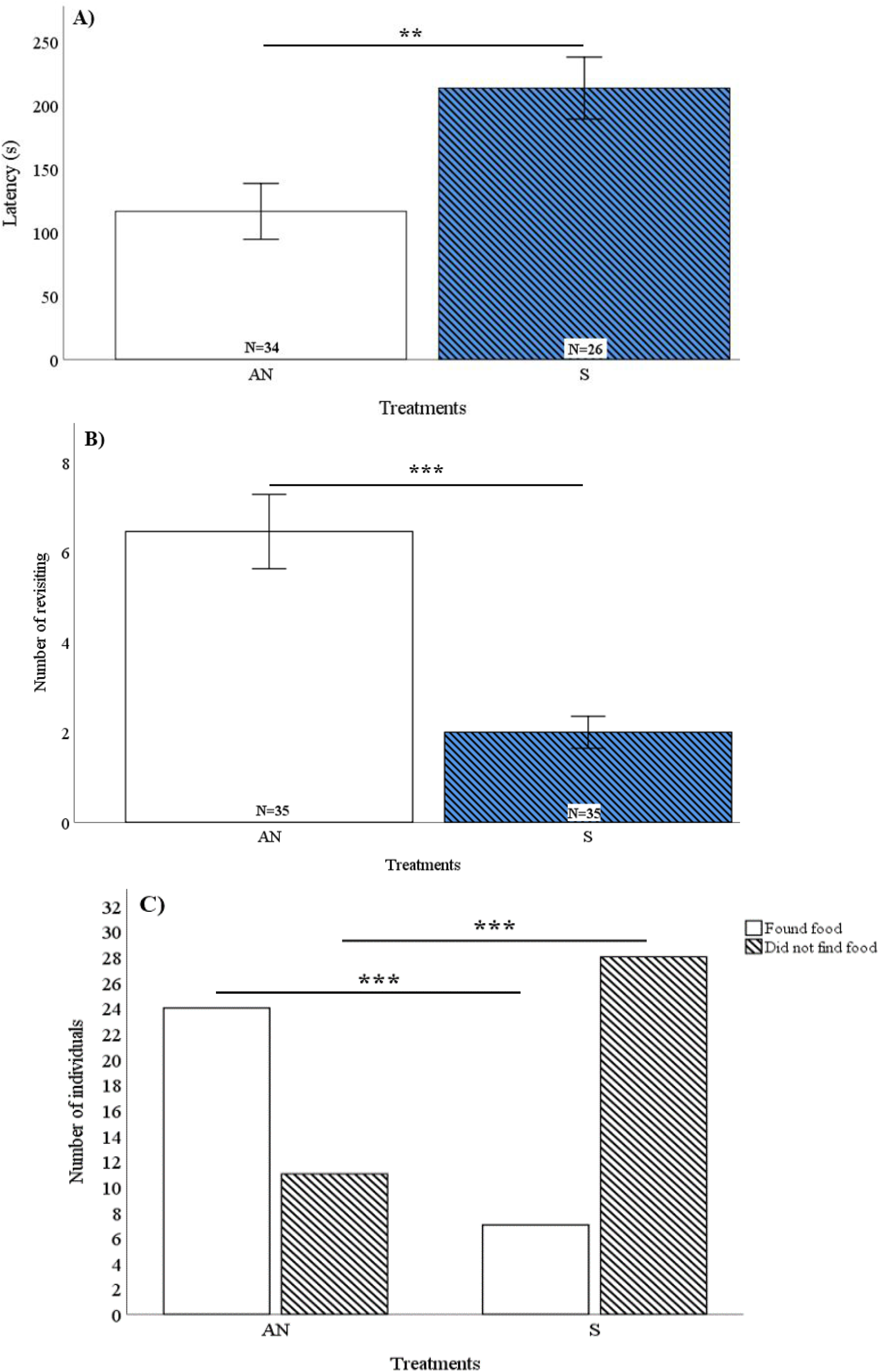
A) Mean ± SE of the time of finding food source in AN: Ambient Noise and S: Sound conditions (N=35). In sound condition the time of finding food significantly increased. B) Mean ± SE of the number of revisiting to food source in AN: Ambient Noise and S: Sound conditions (N=35). In sound condition the number of revisiting to food source significantly decreased. C) The number of individuals that found/not found food source for 10 minutes in AN: Ambient Noise and S: Sound conditions (N=35). In sound condition significantly more shrimp did not find food source. NS: Not Significant, **P<0.01, ***P<0.001.

### The second experiment: feeding distraction

In the sound treatment, more individuals (P<0.001) were distracted during feeding than in the control treatment (Figure 13).

**Fig. 13.**
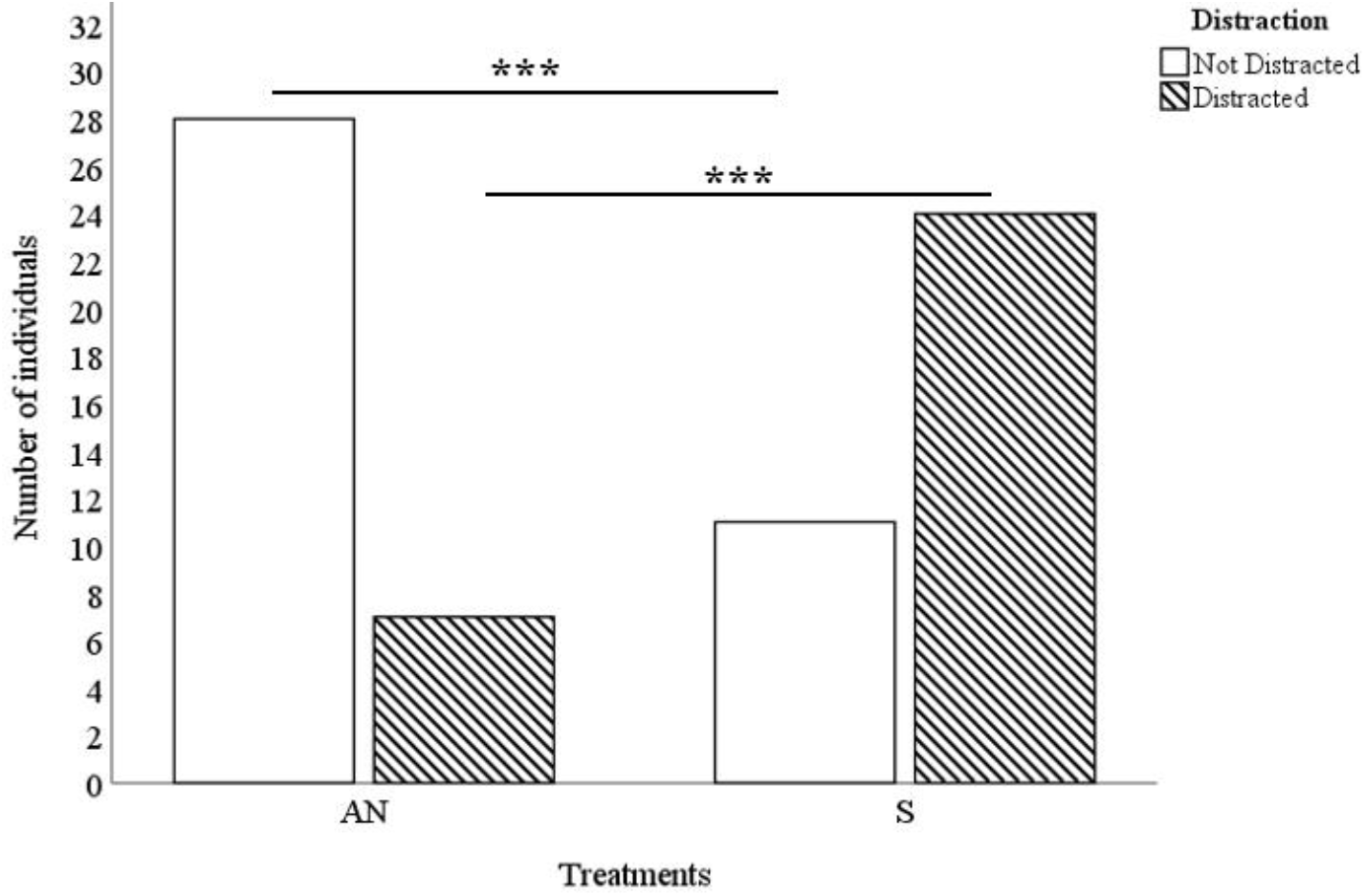
The number of *Neocaridina davidi* that distracted/not distracted from food source for 10 minutes in AN: Ambient Noise and S: Sound conditions (N=35). In sound condition significantly more shrimp distracted in feeding. ***P<0.001.

## Discussion

The impacts of underwater sound on aquatic animals in freshwater habitats have been largely ignored, and the few studies conducted to date show significant impacts on se and fresh-water invertebrates (Breithaupt and Tautz, 1988; Popper et al., 2001; Roberts and Elliott, 2017; Roberts and Laidre, 2019; Ruiz-Ruiz et al., 2020). The direct cascading impacts and indirect consequences of aquatic invertebrates on food domains are quite diverse and important; howeverit is not clear how anthropogenic sound may affect these processes.

The present study is part of a series analyzing the impact of underwater anthropogenic sound on fish and crustaceans (Shafiei Sabet et al., 2020; Mohsenpour and Shafiei Sabet, 2021). This study tested the behavioural effects of acoustic stimuli on the red cherry shrimp, *N. davidi*, which, despite its important ecological and commercial roles, has been poorly investigated in this respect. The present study highlights that invertebrates are likely to perceive sound via their mechanoreceptor organs and be susceptible to the impact of sound as a source of stress. This makes them important candidates for further investigations into the effects of anthropogenic noise pollution.

Experimental sound exposure resulted in a variety of behavioural changes in the red cherry shrimp, some of which have been reported in other invertebrates and some appear to be species-specific. Our results are the first to demonstrate that freshwater decapods can be affected by acoustic stimuli. Sound treatment compromised movement and feeding activity and this is consistent with previous research in crustaceans (Filiciotto et al., 2014; Sal Moyano et al., 2021). In controlled tank-based experiments, it has been shown that sound affects the behaviour of feeding crabs and disrupts their feeding performance (Wale et al., 2013). Moreover, in other taxa, recent studies have shown that acoustic stimuli disrupted feeding performance (Veollmy et al., 2014, Purser and Radford, 2011, Shafiei Sabet et al., 2015; Magnhagen et al., 2017, Mickle and Higgs, 2018).

### First experiment

#### Impact of sound on displacement and locomotion behaviour of shrimp

Upon opening the divider, movement speed decreased in both the control and sound treatments; however, there was no difference between total movement speed for the control and sound treatments. This change may be due to the stress of facing the new environment as shrimps in both treatments displayed the same pattern of reducing their movement speed (Shafiei Sabet et al., 2019). The effects of anthropogenic sound on movement and swimming behaviour of crustaceans and fish is well reported (De Vincenzi et al., 2015, Filiciotto et al., 2016, Sal Moyano et al., 2021, Sarà et al., 2007, Shafiei Sabet 2016a, b, Jimenez et al., 2020). Roberts et al. (2016) found that exposure to anthropogenic substrate-born vibrations, similar to blasting and pile driving, reduced locomotion in adult hermit crabs. Solan et al., (2016), also reported a reduction of locomotor activity when the Norwegian lobster was exposed to continuous and impulsive sound. Snitman et al., (2022) reported that the crab species, *Neohelice granulate*, had reduced locomotion activity when exposed to diverse sound. The reduced locomotion and movement speed could be related to a distraction or confusion effect. It has been shown that Caribbean hermit crabs in response to motor boat sounds were more inactive and allowed a simulated predator to approach closer before their retracting movement. The crabs reallocated finite attention to the distracting acoustic stimuli and were prevented from responding to an approaching threat (Chan et al., 2010). In earthworms, it has been shown that vibrational sounds mask the vibrational cues of approaching foraging moles, making earthworms more prone to predation risks in ensonified areas (Dominoni et al., 2020). Moreover, earthworms may not be able to discriminate between the subterranean waves from anthropogenic sources and vibrations coming from approaching predators (Dominoni et al., 2020). Thus, sound pollution can potentially have multiple impacts on locomotor activities and threat assessment in invertebrates. Other invertebrate studies have shown that acoustic stimuli can have multiple negative effects (Edmonds et al., 2016; Carroll et al., 2017). Finding a mate or detecting a predator has direct effects on reproductive success and survival in animals (Dall et al., 2005; Dominoni et al., 2020). If shrimps spend more of their time budget finding food or shift their attention from foraging areas because of anthropogenic sound, it may detrimentally alter other important biological activities such as courtship and finding a mate to increase their reproductive success or detecting approaching predators to survive in their habitats. These impacts can induce alterations not only on individuals but also can affect populations and ecosystems, especially when anthropogenic sound impacts keynote predators and functionally important habitat forming species. This disruption may have knock-on effects on important ecological and biological processes such as species abundance, survivorship, and the reproductive success of aquatic organisms (Slabbekoorn et al., 2010; Williams et al., 2015; Shafiei Sabet et al., 2016 a,b).

In the control treatment, no difference was observed in the horizontal spatial distribution between individuals, but in the sound treatment, the horizontal distribution of shrimp significantly increased when comparing the point nearest the underwater sound source with the furthest away. This spatial displacement due to sound exposure may have potentially negative impacts on the abundance of invertebrates (Velilla et al., 2021). The authors reported that earthworm abundance decreasedwith increasing vibratory noise. Similar behavior has been reported in other species that have been exposed to annoying or disturbing sounds (De Vincenzi et al., 2021, Shafiei Sabet et al., 2016a, b). In mobile species such as the red cherry shrimp, locomotor movement is an important link between the behaviour of individuals and ecological processes (Herrnkind, 1983; Spanier et al., 1988; Lawton and Lavalli, 1995). Therefore, any impacts of anthropogenic sound on their locomotor activities may have detrimental consequences on their other biological activities such as foraging or anti-predator behaviour. Density-dependent mechanisms underlying the structure and dynamics of populations and communities are often sensitive to short-term changes in the spatial distribution of individuals due to movement (Milinski and Parker, 1991). De Vincenzi et al., (2021) showed that the small-spotted cat shark, when exposed to anthropogenic noise, made different use of the left (close to the underwater speaker) versus the right (away from the underwater speaker) sections of tank, and tended to make different use of the left versus right. Although, the impact of acoustic stimuli on the spatial distribution of red cheery shrimps is not fully understood, in other species there are indications that anthropogenic sound affects swimming activities and spatial displacement. Shafiei Sabet et al., (2016a) reported that sound treatment caused various behavioral changes in both the spatial distribution and swimming behavior of zebrafish within the treatment tank. Sound exposure led to more freezing and less time spent near the active speaker. However, this behavioural response to sound exposure contrasts what was observed in another study on zebrafish under laboratory conditions (Neo et al., 2015). Shafiei Sabet et al., (2016b) compared zebrafishand Lake Victoria cichlids, the former being sensitive to lower absolute thresholds and wider spectral ranges. Consistent with present study, experimental sound exposure induced a significant reduction in swimming speed in the first minute of exposure for both species under captive conditions. Neither species showed spatial shifts away from the active speaker in the horizontal plane. Furthermore, zebrafish showed clear startle response behaviour with the onset of the sound exposure leading to an initial, brief increase in swimming speed, which was not found for the cichlids. In the present study, the experimental set up was successful and able to explore direct assessment of the effects of acoustic stimuli on the horizontal movement of the shrimp. However, it is recognized that parts of the results and behavioural observations may suffer from the bias introduced by experimental and acoustics artifacts (Rogers et al., 2016; Parvulescu, 1967; Akamatsu et al., 2002). Particle motion components of the experimental set up were not able to be measured, which makes it likely that the behavioural observations were in response to particle motion component rather than sound pressure (Popper and Hawkins, 2018; Hawkins et al., 2021).

#### Acoustic stimuli compromised feeding performance in shrimp

Anthropogenic sounds were found to have detectable effects on the foraging performance of fish and invertebrates. In the present study, feeding latency in shrimps significantly increased in the sound treatment compared to the control. This delay in finding the food source seems to be due to stress and confusion caused by sound playback, which agrees with previous studies using crustaceans and fish (Hughes et al., 2014, Gendron et al., 2020, Voellmy et al., 2013). Elevated levels of sound in the present study had negative effects on the feeding activity and performance in shrimp under laboratory conditions.

More individuals successfully found the food source in the control treatment than the sound treatment, which is consistent with the findings of Voellmy et al. (2013) and confirms the effect of sound on feeding distraction. These observations suggest that anthropogenic sound may cause similar impacts on foraging activities in animals under laboratory conditions. Although it is important to consider that the findings of the present study should not be interpreted for wild animals and fish in field conditions. Various studies have shown that food-deprivation influences behaviour including foraging; food-deprived three-spined sticklebacks were more likely to initiate predator inspection visits and had higher feeding rates than well-fed shoal mates (Godin and Crossman, 1994). Hungry fifteen-spined sticklebackswere more likely to feed at a food source associated with a predator threat (and therefore, were less risk-averse) than partially satiated individuals (Croy and Hughes, 1991).

The number of revisits to the food source also significantly decreased when exposed to sound. Such changes in feeding behavior have not been reported previously. This study showed that anthropogenic noise pollution influences the behavioural activity of the red cherry shrimp. Currently, no data on the sensitivity of red cherry shrimp to acoustic signals is available; however, the results of the present study indicate that *N. davidi* may perceive all or part of the acoustic stimuli projected within the wider band-width of their underwater audible rasps characterized by signals with most of energy concentrated in the 4-2000 Hz range, with the peak frequency of 700 Hz and peak amplitude of 120 dB re 1 lPa (Buscaino et al., 2011b). Filiciotto et al., (2014) found that lobsters exposed to boat sounds had higher mobility and moving state values compared to lobsters in the control treatment. Shafiei Sabet et al., (2019) found no significant changes in behavioural parameters of *Daphnia magna* while Sal Moyano et al. (2021) evaluated the effects of biological and anthropogenic acoustic signals on the orientation response of different stages (megalopae and juveniles) of 4 brachyuran crabs species. *C. angulatus* megalopae and juveniles responded positively towards crustacean signals, while juveniles responded negatively towards fish sounds. *N. granulata* juveniles orientated negatively towards crustacean, motorboat and fish signals while *C. altimanus* and *L. uruguayensis* juveniles did not respond to fish signals (Sal Moyano et al., 2021). The results support the idea that invertebrates can discriminate among conspecific signals and highlight the role of sound on prey-predator relationships. The behavioral orientation response to the motorboat sound demonstrates a presumably negative effect of anthropogenic sound on the biological interactions of species.

There is mounting experimental evidence that anthropogenic sound can have a variety of negative physiological and behavioural effects on individual animals, ultimately affecting their survival and reproductive success (Kight and Swaddle, 2011; Shannon et al., 2015; Kunc et al., 2016; Morley et al., 2014). For example, sound from human activities can directly cause injury or hearing loss in some species (Halvorsen et al., 2012; Popper et al., 2005),. It can also mask biologically important sounds, such as the sounds the fish make themselves or the sound of predators (i.e. reducing the signal-to-noise ratio). It reduces the range over which fish can detect biologically important sounds and subsequently impose them to negative fitness output (Popper et al., 2005). Elevated ambient sound can also impair the ability of animals to communicate (Slabbekoorn and Ripmeester, 2008, Kunc et al., 2015; Lampe et al., 2012). Sound can be perceived as a threat, a distraction, or can cause increased stress, in turn impairing an animal’s ability to forage efficiently (Wale et al., 2013; Voellmy et al., 2014). It can also impair an animal’s ability to respond appropriately to information about predation risk (Morris-Drake et al., 2016; Simpson et al., 2016; Purser et al., 2016), perform adaptive behaviours during habitat-selection (Holles et al., 2013; Simpson et al., 2016), or reproduce successfully (Blickley et al., 2012; Nedelec, 2017).

Studies on other taxa have revealed that sound affects swimming activities and spatial distribution. Filiciotto et al., (2016) found that the common prawn had clear behavioural responses to acoustic stimuli exposure (boat noise). In particular, those exposed to sounds had significantly lower encounters between subjects but spent more time outside the shelter and resting. Sound exposure caused behavioural modifications and negative effects on feeding frequencies in the mediterranean damselfish (Bracciali et al., 2012). Sarà et al., (2007) found that in the absence of boat noise, tuna assumed a concentrated coordinated school structure with unidirectional swimming and without a precise shape. When a car ferry approached, tuna changed swimming direction and increased their vertical movement towards the surface or bottom. Theschool exhibited an unconcentrated structure and uncoordinated swimming behaviour. Shafiei Sabet et al., (2016b) compared zebrafish and Lake Victoria cichlids, the former being sensitive to lower absolute thresholds and wider spectral ranges. Experimental sound exposure caused a significant reduction in swimming speed in the first minute of exposure for both species under captive conditions. Furthermore, zebrafish showed clear startle response behaviour with the onset of the sound exposure leading to an initial, brief increase in swimming speed, which was not found for the cichlids. Neither species showed spatial shifts away from the active speaker in the horizontal plane. Blom et al., (2019) tested the impact of broadband sound exposure (added either continuously or intermittently)on the behaviour and reproductive success of the common goby. Compared to the intermittent noise and control treatments, the continuous noise treatment increased latency to female nest inspection.

### Second experiment

#### Does an acoustic stimulus compromise food source assessment and attention in shrimp?

In the sound treatment, significantly more individuals were distracted in feeding compared to the control treatment. Similar results are reported in other species (Wale et al., 2013, Hastie et al., 2021, Gendron et al., 2020, Hughes et al., 2014, Voellmy et al., 2013). Wale et al., (2013) argued that crabs exposed to ship noise were more likely to suspend feeding than those exposed to ambient-noise. Gendron et al., (2020) found the presence of boat noise had a significant effect on the hunting behaviour of *Pseudopleuronectes americanus* larvae. Larvae exposed to boat noise spent less time hunting and had smaller stomach volumes compared to those exposed to no sound. This suggests that more prey were consumed in the absence of boat noise. Purser and Radford (2011) noted a decrease in foraging performance in three-spined sticklebacks exposed to noise. This decrease resulted from (1) the misidentification of food versus non-food items, as shown by an increased number of attacks on the latter, and (2) fewer successful attacks on food items under noisy conditions. Hanache et al., (2018) reported that boat noise significantly reduced attack rate, resulting in a functional response curve of the same height but with a less steep initial slope. European minnows exhibited a stress response to noise including increased swimming distance, more social interactions, and altered spatial distribution. McLaughlin and Kunc., (2015) found that cichlids exposed to anthropogenic noise showed an increase in sheltering accompanied by a decrease in foraging. Their results highlight the multiple negative effects of an environmental stressor on an individual’s behaviour. In an experiment where crabs could only find food based on olfaction, Hubert et al., (2021) found no overall effect of boat sound on food finding success and efficacy, foraging duration or walking distance in shore crabs. There are several mechanisms to explain how anthropogenic sound affects behaviour: increasing stress (Smith et al., 2004), masking biological relevant cues (Hawkins and Chapman, 1975), distracting of attention (Chan et al., 2010; Dukas, 2004) and physically damaging sensory system and internal organs (McCauley et al., 2003). These could be operating separately or simultaneously, and in many cases, it is difficult to determine which of these is most relevant, especially in the present study.

Wale et al., (2013) showed that the ability of crabs to find food items was not impaired by the playback of ship noise compared to ambient noise. Individuals exposed to ship noise were more likely to suspend feeding than those exposed to ambient-noise. Hughes et al., (2014) showed that the sounds made by predatory fish can affect the feeding behavior of crabs. In the presence of the sound of predatory fish in the frequency range of 10 to 1600 Hz and the sound intensity of 146 to 116 dB, the amount of crab feeding from bivalves was significantly reduced.

Sound exposure can affect feeding activities in other taxa. Hastie et al (2021) found that sound can negatively affect the foraging activity of gray seals, with the success of seal foraging being higher in the control treatment than the sound treatment. Goldbogen et al (2013) demonstrated that sound can significantly affect blue whales during deep feeding behaviour. These sound related effects on foraging activities that have been observed in other species agree with the results of the present study.

Although it needs further research, shrimps that do not feed properly because of acute and/or chronic exposure to sound could be affected in their growth, size at sexual maturity, mate selection and brooding eggs behavior, thus affecting the whole life cycle and the species fitness and this role in the trophic webs. Sounds generated by human activities are interfering with biologically relevant acoustic information of aquatic animals/species across taxa with important population and community level consequences (Derryberry et al., 2020; Dominoni et al., 2020).

Our results provide important insights into the effects of sound on this species at individual level under laboratory conditions. Opening the divider caused a non-auditory behavioural response by reducing the instant movement of shrimps, which may be an anti-predator response and likely improves their survival by lowering visual visibility. Shrimps spent more time away from the active speaker suggesting that they were able to detect the acoustic stimuli and further research measuring both sound pressure and particle motion components is recommended. Anthropogenic sound decreased food finding success and the number of revisits to the food source, increased feeding latency and distraction. Overall, the findings of the present study highlight the negative impacts of anthropogenic sound on the behaviour of shrimps that could impair their foraging performance and potentially have cascading effects on their survival.

## Conclusions

It was observed how short-term exposure to sound changes some behavioural patterns in *N. davidi*. The simulated sound in laboratory conditions influenced the behaviour of specimens, mainly in terms of spatial distribution, feeding latency, finding-food and number of revisits to the food source and feeding distraction. As new information becomes available, there is a need to change the sound exposure criteria and implement updates of invertebrates’ perception with regard to the acoustic environment. The metrics used currently are often inappropriate as they are expressed in terms of sound pressure and potential impacts on invertebrates. Whereas, all fishes and most invertebrates are able to perceive and detect particle motion components (Popper and Hawkins, 2019; Hawkins et al., 2021). There is a need to explore anthropogenic sound effects on invertebrates both in terms of sound pressure levels and particle motion components.

The actual impacts of sound on populations of invertebrates and other taxa are often unknown and difficult to access. Longer-term studies under laboratory conditions are necessary to explore the potential chronic effects of sound on populations and on ecological communities reverberating among food webs. However, it is unknown what impact this will have on much larger populations of animals in terms of sound-related impacts and ecological effects. Future experiments should be performed in order to assess the animal responses in nature where the acoustic field is not influenced by laboratory conditions. Moreover, long-term field studies should be carried out to evaluate the effects of noise from human activity on some ecological and conservation factors in this species, providing and implementing innovative tools for the measure of the acoustic pollution phenomenon.

## References

Akamatsu, T., Okumura, T., Novarini, N., Yan, H.Y., 2002. Empirical refinements applicable to the recording of fish sounds in small tanks. J. Acoust. Soc. Am. 112, 3073e3082. http://dx.doi.org/10.1121/1.1515799.

Andrew, J., Linda, A., Wright, A. J., Beale, C. M., Martineau, D., 2007. Do Marine Mammals Experience Stress Related to Anthropogenic Noise?, Inter. J. Comp. Psycho. 20, 274–316.

Andrew, R. K., Howe, B. M., Mercer, J. A., 2011. Long-time trends in ship traffic noise for four sites off the North American West Coast. J. Acoust. Soc. Am. 129(2), 642. https://doi.org/10.1121/1.3518770.

ASAB, 2020. Guidelines for the treatment of animals in behavioural research and teaching. Anim. Behav. 135. I–X.

Bell, S. S., Coull, B. C., 1978. Field evidence that shrimp predation regulates meiofauna. Oecologia. 35(2), 141–148. https://doi.org/10.1007/BF00344727.

Blickley, J. L., Blackwood, D., Patricelli, G. L., 2012. Experimental Evidence for the Effects of Chronic Anthropogenic Noise on Abundance of Greater Sage-Grouse at Leks. Conserv. Biol, 26(3), 461–471. https://doi.org/10.1111/J.1523-1739.2012.01840.X.

Blom, E. L., Kvarnemo, C., Dekhla, I., Schöld, S., Andersson, M. H., Svensson, O., Amorim, M. C. P., 2019. Continuous but not intermittent noise has a negative impact on mating success in a marine fish with paternal care. Sci. Rep. 9(1), 1–9. https://doi.org/10.1038/s41598-019-41786-x.

Bracciali, C., Campobello, D., Giacoma, C., Sarà, G., 2012. Effects of Nautical Traffic and Noise on Foraging Patterns of Mediterranean Damselfish (*Chromis chromis*). PLoS ONE. 7(7), 40582. https://doi.org/10.1371/JOURNAL.PONE.0040582.

Breithaupt, T., & Tautz, J., 1988. Vibration sensitivity of the crayfish statocyst. Naturwissenschaften. 75, 310–312.

Bruintjes, R., Purser, J., Everley, K. A., Mangan, S., Simpson, S. D., Radford, A. N., 2015. Rapid recovery following short-term acoustic disturbance in two fish species. Roy. Soc. Op. Sci. 3(1). https://doi.org/10.1098/RSOS.150686.

Brumm, H., & Slabbekoorn, H., 2005. Acoustic Communication in Noise. Adv. Stu. Behav. 35, 151–209. https://doi.org/10.1016/S0065-3454(05)35004-2.

Buscaino, G., Filiciotto, F., Gristina, M., Buffa, G., Bellante, A., Maccarrone, V., Patti, B., Mazzola, S., 2011. Defensive strategies of European spiny lobster *Palinurus elephas* during predator attack. Mar. Ecol. Prog. 423, 143–154.

Carroll, A. G., Przeslawski, R., Duncan, A., Gunning, M., Bruce, B., 2017. A critical review of the potential impacts of marine seismic surveys on fish & invertebrates. Mar. Poll. Bull. 114(1), 9–24. https://doi.org/10.1016/j.marpolbul.2016.11.038.

Chan, A. A. Y. H., Blumstein, D. T., 2011. Attention, noise, and implications for wildlife conservation and management. App. Anim. Behav. Sci. 131(1-2), 1–7. https://doi.org/10.1016/J.APPLANIM.2011.01.007.

Chan, A. A. Y. H., Giraldo-Perez, P., Smith, S., Blumstein, D. T., 2010. Anthropogenic noise affects risk assessment and attention: the distracted prey hypothesis. Biol. Lett. 6(4), 458–461. https://doi.org/10.1098/RSBL.2009.1081.

Charmandari, E., Tsigos, C., Chrousos, G., 2004. Endocrinology of the stress response. Annu. Rev. Physiol. 67, 259–284. https://doi.org/10.1146/ANNUREV.PHYSIOL.67.040403.120816.

Cousteau, J., Dumas, F., 1987. The Silent World. N. Lyons Books, New York, NY.

Croy, M.I., Hughes, R.N., 1991. Effects of food supply, hunger, danger and competition on choice of foraging location by the fifteen-spined stickleback, *Spinachia spinachia* L. Anim. Behav. 42, 131–139.

Dall, S. R. X., Giraldeau, L. A., Olsson, O., McNamara, J. M., & Stephens, D. W., 2005. Information and its use by animals in evolutionary ecology. Trends Ecol. Evol. 20(4), 187–193. https://doi.org/10.1016/J.TREE.2005.01.010.

De Jong, K., Amorim, M. C. P., Fonseca, P. J., Fox, C. J., Heubel, K. U., 2018. Noise can affect acoustic communication and subsequent spawning success in fish. Environ. Pollut. 237, 814–823. https://doi.org/10.1016/J.ENVPOL.2017.11.003.

De Kloet, E. R., Oitzl, M. S., Joëls, M., 1999. Stress and cognition: are corticosteroids good or bad guys?. Trends Neurosci. 22(10), 422–426. https://doi.org/10.1016/S0166-2236(99)01438-1.

De Vincenzi, G., Maccarrone, V., Filiciotto, F., Buscaino, G., Mazzola, S., 2015. Behavioural responses of the European Spiny lobster, *Palinurus elephas* (Fabricius, 1787), to conspecific and synthetic sounds. Crustaceana. 88(5), 523–540.https://doi.org/10.1163/15685403-00003430.

Derryberry, E. P., Phillips, J. N., Derryberry, G. E., Blum, M. J., Luther, D., 2020. Singing in a silent spring: Birds respond to a half-century soundscape reversion during the COVID-19 shutdown. Science. 370(6516), 575–579.

Dominoni, D. M., Halfwerk, W., Baird, E., Buxton, R. T., Fernández-Juricic, E., Fristrup, K. M., McKenna, M. F., Mennitt, D. J., Perkin, E. K., Seymoure, B. M., Stoner, D. C., Tennessen, J. B., Toth, C. A., Tyrrell, L. P., Wilson, A., Francis, C. D., Carter, N. H., Barber, J. R., 2020. Why conservation biology can benefit from sensory ecology. Nat. Ecol. Evol. 4(4), 502–511. https://doi.org/10.1038/S41559-020-1135-4.

Duarte, C. M., Chapuis, L., Collin, S. P., Costa, D. P., Devassy, R. P., Eguiluz, V. M., Erbe, C., Gordon, T. A. C., Halpern, B. S., Harding, H. R., Havlik, M. N., Meekan, M., Merchant, N. D., Miksis-Olds, J. L., Parsons, M., Predragovic, M., Radford, A. N., Radford, C. A., Simpson, S. D., Juanes, F., 2021. Soundscape Anthro. ocean. Sci. 371(6529). https://doi.org/10.1126/SCIENCE.ABA4658/SUPPL_FILE/ABA4658_MDAR_REPRODUCIBILITY_CHECKLIST.PDF.

Dukas, R., 2004. Causes and consequences of limited attention. Brain. Behav. Evol. 63(4): 197–210.

Edmonds, N. J., Firmin, C. J., Goldsmith, D., Faulkner, R. C., Wood, D. T., 2016. A review of crustacean sensitivity to high amplitude underwater noise: Data needs for effective risk assessment in relation to UK commercial species. Mar. Pollut. Bull. 108(1-2),5–11. https://doi.org/10.1016/J.MARPOLBUL.2016.05.006.

Filiciotto, F., Vazzana, M., Celi, M., Maccarrone, V., Ceraulo, M., Buffa, G., Stefano, V. Di, Mazzola, S., Buscaino, G., 2014. Behavioural and biochemical stress responses of *Palinurus elephas* after exposure to boat noise pollution in tank. Mar. Pollut. Bull. 84(1-2), 104–114. https://doi.org/10.1016/j.marpolbul.2014.05.029.

Filiciotto, F., Vazzana, M., Celi, M., Maccarrone, V., Ceraulo, M., Buffa, G., Arizza, V., de Vincenzi, G., Grammauta, R., Mazzola, S., Buscaino, G., 2016. Underwater noise from boats: Measurement of its influence on the behaviour and biochemistry of the common prawn (*Palaemon serratus*, Pennant 1777). J. Exp. Mar. Biol. 478, 24–33. https://doi.org/10.1016/jjembe.2016.01.014.

Frisk, G. V., 2012. Noiseonomics: The relationship between ambient noise levels in the sea and global economic trends. Sci. Rep. 2(1), 1–4.https://doi.org/10.1038/srep00437.

Gaston, K. J., Visser, M. E., Hölker, F., 2015. The biological impacts of artificial light at night: The research challenge. Philos. Trans. R. Soc. B: Biological Sciences. 370(1667). https://doi.org/10.1098/RSTB.2014.0133.

Gendron, G., Tremblay, R., Jolivet, A., Olivier, F., Chauvaud, L., Winkler, G., Audet, C., 2020. Anthropogenic boat noise reduces feeding success in winter flounder larvae (*Pseudopleuronectes americanus*). Environ. Biol. Fishes. 103(9), 1079–1090. https://doi.org/10.1007/s10641-020-01005-3.

Godin, J.J., Crossman, S.L., 1994. Hunger-dependent predator inspection and foraging behaviours in the three spine stickleback (*Gasterosteus aculeatus*) under predation risk. Behav. Ecol. Sociobiol. 34, 359–366

Goldbogen, J. A., Southall, B. L., DeRuiter, S. L., Calambokidis, J., Friedlaender, A. S., Hazen, E. L., Tyack, P. L., 2013. Blue whales respond to simulated mid-frequency military sonar. Proc. Royal Soc. B: Biological Sciences, 280(1765), 20130657.

Gordon, J., Tyack, P.L., 2001. Marine Mammals: Biology and Conservation. Kjuwer Academic/Plenum, 2002.

Halvorsen, M. B., Casper, B. M., Matthews, F., Carlson, T. J., Popper, A. N., 2012. Effects of exposure to pile-driving sounds on the lake sturgeon, Nile tilapia and hogchoker. Proc. R. Soci. B: Biological Sciences, 279(1748), 4705–4714. https://doi.org/10.1098/RSPB.2012.1544.

Hanache, P., Spataro, T., Firmat, C., Boyer, N., Fonseca, P., Médoc, V., 2020. Noise-induced reduction in the attack rate of a planktivorous freshwater fish revealed by functional response analysis. Freshw. Biol. 65(1), 75–85. https://doi.org/10.1111/fwb.13271.

Hastie, G. D., Lepper, P., McKnight, J. C., Milne, R., Russell, D. J. F., Thompson, D., 2021. Acoustic risk balancing by marine mammals: anthropogenic noise can influence the foraging decisions by seals. J. Appl. Ecol. 58(9), 1854–1863. https://doi.org/10.1111/1365-2664.1391

Hawkins, A. D., Myrberg, A. A., 1983. Hearing and sound communication under water. Bioacoustics, a comparative approach. Academic Press, London, 347–405.

Hawkins, A. D., Hazelwood, R. A., Popper, A. N., Macey, P. C., 2021. Substrate vibrations and their potential effects upon fishes and invertebrates. J. Acoust. Soc. Am. 149(4), 2782–2790. https://doi.org/10.1121/10.0004773.

Hawkins, A. D., Roberts, L., Cheesman, S., 2014. Responses of free-living coastal pelagic fish to impulsive sounds. J. Acoust. Soc. Am. 135(5), 3101–3116.https://doi.org/10.1121/1.4870697.

Hawkins, A.D., Chapman, C.J., 1975. Masked auditory thresholds in the cod, *Gadus morhua*. J. Comp. Physiol. A, 103, 209–226.

Herbert-Read, J. E., Kremer, L., Bruintjes, R., Radford, A. N., Ioannou, C. C., 2017. Anthropogenic noise pollution from pile-driving disrupts the structure and dynamics of fish shoals. Proc. R. Soc. B: Biol. Sci. 284(1863). https://doi.org/10.1098/RSPB.2017.1627

Herrnkind, W. F., 1983. Movements Patterns and Orientation. In: F. J. Vernberg W. B. Vernberg (Eds.), The Biology of Crustacea, Vol. 7. Behav. Ecol. 41–105. (Academic Press, New York, Ny).

Hildebrand, J., 2009. Anthropogenic and natural sources of ambient noise in the ocean. Mar. Ecol. Prog. Ser. 395, 5–20. https://doi.org/10.3354/meps08353.

Holles, S. H., Simpson, S. D., Radford, A. N., Berten, L., Lecchini, D., 2013. Boat noise disrupts orientation behaviour in a coral reef fish. Mar. Ecol. Prog. Ser. 485, 295–300. https://doi.org/10.3354/MEPS10346.

Hubert, J., van Bemmelen, J. J., Slabbekoorn, H., 2021. No negative effects of boat sound playbacks on olfactory-mediated food finding behaviour of shore crabs in a T-maze. Environ. Pollut. 270, 116184. https://doi.org/10.1016/j.envpol.2020.116184.

Hughes, A., Mann, D. A., Kimbro, D. L., 2014. Predatory fish sounds can alter crab foraging behaviour and influence bivalve abundance. Proc. R. Soc. B: Biological Sciences, 281(1788). https://doi.org/10.1098/rspb.2014.071.

Jimenez, V.L., Fakan, E. P., McCormick, M. I., 2020. Vessel noise affects routine swimming and escape response of a coral reef fish. PloS One, 15(7), e0235742. https://doi.org/10.1371/journal.pone.0235742.

Kelly, E., Tully, O., Browne, R., 2012. Effects of temperature and salinity on the survival and development of larval and juvenile *Palaemon serratus* (Decapoda: Palaemonidae) from Irish waters. J. Mar. Biolog. Assoc. United Kingdom, 92(1), 151–161. https://doi.org/10.1017/S0025315411000415.

Kight, C. R., Swaddle, J. P., 2011. How and why environmental noise impacts animals: An integrative, mechanistic review. Ecol. Lett. 14(10), 1052–1061. https://doi.org/10.1111/J.1461-0248.2011.01664.X.

Klotz, W., Miesen, F. W., Hüllen, S., & Herder, F., 2013. Two Asian fresh water shrimp species found in a thermally polluted stream system in north Rhine-Westphalia, Germany. Aquat. Invasions. 8(3), 333–339. https://doi.org/10.3391/AI.2013.8.3.09.

Kunc, H. P., Lyons, G. N., Sigwart, J. D., McLaughlin, K. E., Houghton, J. D. R., 2015. Anthropogenic Noise Affects Behavior across Sensory Modalities. Am. Nat. https://Doi.Org/10.1086/677545, 184(4), E93–E100. https://doi.org/10.1086/677545.

Kunc, H. P., McLaughlin, K. E., Schmidt, R., 2016. Aquatic noise pollution: implications for individuals, populations, and ecosystems. Proc. R. Soc. B: Biological Sciences. 283(1836). https://doi.org/10.1098/RSPB.2016.0839.

Lampe, U., Schmoll, T., Franzke, A., Reinhold, K., 2012. Staying tuned: grasshoppers from noisy roadside habitats produce courtship signals with elevated frequency components. Funct. Ecol. 26(6), 1348–1354.https://doi.org/10.1111/1365-2435.12000.

Lara, R. A., Vasconcelos, R. O., 2019. Characterization of the Natural Soundscape of Zebrafish and Comparison with the Captive Noise Conditions. Zebrafish. 16(2), 152–164. https://doi.org/10.1089/zeb.2018.1654.

Lawton, P. K. L,. Lavalli, N., 1995. Postlarval, Juvenile, Adolescent, and Adult Ecology. In: J. R. Factor (Ed.), Biology of the Lobster Homarus americanus: 47–88. (Academic Press, San Diego, Ca).

Liang, X.Q., 2004. Fauna Sinica. Invertebrata-Crustacea-Decapoda-Atyidae.Science Press, Beijing, China.

Lopeztegui-Castillo, A., 2021. Assessment of nutritional condition in crustaceans: a review of methodologies and guidelines for applying inexpensive and wide-ranging indices to the spiny lobster *Panulirus argus* (Latreille, 1804) (Decapoda: Achelata: Palinuridae). J. Crustac. Biol. 41(4). https://doi.org/10.1093/JCBIOL/RUAB067.

Lovell, J. M., Findlay, M. M., Moate, R. M., Yan, H. Y., 2005. The hearing abilities of the prawn *Palaemon serratus*. Comp. Biochem. Physiol. A Mol. Integr. Physiol. 140(1), 89–100. https://doi.org/10.1016/J.CBPB.2004.11.003.

Lupien, S. J., McEwen, B. S., 1997. The acute effects of corticosteroids on cognition: integration of animal and human model studies. Brain Res. Rev. 24(1), 1–27.https://doi.org/10.1016/S0165-0173(97)00004-0.

Magnhagen, C., Johansson, K., Sigray, P., 2017. Effects of motorboat noise on foraging behaviour in Eurasian perch and roach: A field experiment. Mar. Ecol. Prog. Ser. 564, 115–125. https://doi.org/10.3354/meps1197.

McCauley, R.D., Fewtrell, J., Popper, A.N., 2003. High intensity anthropogenic sound damages fish ears. J. Acoust. Soc. Am. 113:1–5

McDonald, M. A., Hildebrand, J. A., Wiggins, S. M., 2006. Increases in deep ocean ambient noise in the Northeast Pacific west of San Nicolas Island, California J. Acoust. Soc. Am. 120(2), 711. https://doi.org/10.1121/1.2216565.

McLaughlin, K. E., Kunc, H. P. 2015. Changes in the acoustic environment alter the foraging and sheltering behaviour of the cichlid *Amititlania nigrofasciata*. Behav. Processes. 116, 75–79. https://doi.org/10.1016/j.beproc.2015.04.012.

Mendl, M., 1999. Performing under pressure: stress and cognitive function. Appl. Anim. Behav. Sci. 65(3), 221–244. https://doi.org/10.1016/S0168-1591(99)00088-X.

Mickle, M. F., Higgs, D. M., 2018. Integrating techniques: A review of the effects of anthropogenic noise on freshwater fish. Can. J. Fish. Aquat. Sci. 75(9), 1534–1541. https://doi.org/10.1139.

Milinski, M., Parker, G.A., 1991. Competition for resources. In: Krebs, J.R., Davies, N.B. (Eds.), Behavioural Ecology. An Evolutionary Approach. Blackwell Scientific Publications, Oxford, p. 137–168.

Mohsenpour, R., Shafiei Sabet, S., 2021. Spatial distribution of zebrafish (*Danio rerio*) as a behavioral index of stress in response to sound. Aqua. Phys. Biotech. 9(3), 23–46. https://doi.org/10.22124/JAPB.2021.17584.1393.

Morley, E. L., Jones, G., Radford, A. N., 2014. The importance of invertebrates when considering the impacts of anthropogenic noise. Proc. R. Soc. B: Biological Sciences, 281(1776). https://doi.org/10.1098/RSPB.2013.2683.

Morris-Drake, A., Kern, J. M., Radford, A. N., 2016. Cross-modal impacts of anthropogenic noise on information use. Curr. Biol. 26(20), R911–R912. https://doi.org/10.1016/J.CUB.2016.08.064/ATTACHMENT/E79CD989-FEEE-468C-8BB8-3CB5F9514D64/MMC1.PDF.

Nedelec, S. L., Radford, A. N., Pearl, L., Nedelec, B., McCormick, M. I., Meekan, M. G., Simpson, S. D., 2017. Motorboat noise impacts parental behaviour and offspring survival in a reef fish. Proc. R. Soc. B: Biol. Sci. 284(1856). https://doi.org/10.1098/RSPB.2017.0143.

Neo, Y. Y., Parie, L., Bakker, F., Snelderwaard, P., Tudorache, C., Schaaf, M., Slabbekoorn, H., 2015. Behavioral changes in response to sound exposure and no spatial avoidance of noisy conditions in captive zebrafish. Front. Behav. Neurosci. 9, 28.

Nowacek, D. P., Thorne, L. H., Johnston, D. W., Tyack, P. L., 2007. Responses of cetaceans to anthropogenic noise. Mamm. Rev. 37(2), 81–115. https://doi.org/10.1111/J.1365-2907.2007.00104.

Onuki, K., Fuke, Y., 2022. Rediscovery of a native freshwater shrimp, *Neocaridina denticulata*, and expansion of an invasive species in and around Lake Biwa, Japan: genetic and morphological approach. Conserv. Genet. 1–14. https://doi.org/10.1007/S10592-022-01467-1.

Parvulescu, A., 1967. The acoustics of small tanks. In: Tavolga, W.N. (Ed.), Marine Bio-Acoustics. Pergamon Press, Oxford

Payne, R., Webb, D., 1971. Orientation by means of long range acoustic signaling in baleen whales. Annals of the New York Academy of Sciences, 188(1), 110–141.https://doi.org/10.1111/J.1749-6632.1971.TB1309348.

Peng, C., Zhao, X., Lio, G., 2015. Noise in the sea and its impact on marine organisms, International Int. J. Environ. Res. Public Health, 12, 12304–12323.

Popper, A. N., Hawkins, A. D., 2019. An overview of fish bioacoustics and the impacts of anthropogenic sounds on fishes. J. Fish Biol. 94(5), 692–713. https://doi.org/10.1111/JFB.13948.

Popper, A. N., Salmon, M., Horch, K. W., 2001. Acoustic detection and communication by decapod crustaceans. J. Comp. Physiol. A 2001 187:2, 187(2), 83–89. https://doi.org/10.1007/S003590100184.

Popper, A. N., Smith, M. E., Cott, P. A., Hanna, B. W., MacGillivray, A. O., Austin, M. E., Mann, D. A., 2005. Effects of exposure to seismic airgun use on hearing of three fish species. J. Acoust. Soc. Am. 117(6), 3958. https://doi.org/10.1121/1.1904386.

Popper, A.N., 2003. Effects of anthropogenic sounds on fishes. Fisheries, 28, 24–31

Purser, J., Radford, A. N., 2011. Acoustic Noise Induces Attention Shifts and Reduces Foraging Performance in Three-Spined Sticklebacks (*Gasterosteus aculeatus*). PLOS ONE, 6(2), e17478. https://doi.org/10.1371/JOURNAL.PONE.0017478.

Purser, J., Bruintjes, R., Simpson, S. D., Radford, A. N., 2016. Condition-dependent physiological and behavioural responses to anthropogenic noise. Physiol. Behav. 155, 157–161. https://doi.org/10.1016/J.PHYSBEH.2015.12.010.

Quinn, J., J. Whittingham, M. J., Butler, S., Cresswell, W., 2006. Noise, predation risk compensation and vigilance in the chaffinch *Fringilla coelebs*. J. Avian Biol. 37(6), 601–608. https://doi.org/10.1111/J.2006.0908-8857.03781.X.

Radford, C., Slater, M., 2019. Soundscapes in aquaculture systems. Aquac. Environ. Interact. 11, 53–62. https://doi.org/10.3354/aei00293.

RStudio Team, (2020). RStudio: Integrated Development for R. RStudio, PBC, Boston, MA URL http://www.rstudio.com.

Reise, K., 1979. Moderate predation on meiofauna by the macrobenthos of the Wadden Sea. Helgoländer Wissenschaftliche. Meeresuntersuchungen. 32(4), 453–465. https://doi.org/10.1007/BF02277989.

Richardson, W.J., Greene Jr., C.R., Malme, C.I., Thomson, D.H., 1995. Marine Mammals and Noise. Academic Press, New York.

Roberts, L., Elliott, M., 2017. Good or bad vibrations? Impacts of anthropogenic vibration on the marine epibenthos. Sci. Total Environ. 595, 255–268. https://doi.org/10.1016/J.SCITOTENV.2017.03.117.

Roberts, L., Laidre, M. E., 2019. Finding a home in the noise: Cross-modal impact of anthropogenic vibration on animal search behaviour. Biol. Open. 8(7). https://doi.org/10.1242/BIO.041988/2242.

Roberts, L., Cheesman, S., Breithaupt, T., Elliott, M., 2015. Sensitivity of the mussel *Mytilus edulis* to substrate-borne vibration in relation to anthropogenically generated noise. Mar. Ecol. Prog. Ser. 538, 185–195.https://doi.org/10.3354/MEPS11468.

Roberts, L., Cheesman, S., Elliott, M., Breithaupt, T., 2016. Sensitivity of *Pagurus bernhardus* (L.) to substrate-borne vibration and anthropogenic noise. J. Exp. Mar. Biol. 474, 185–194. https://doi.org/10.1016/jjembe.2015.09.014.

Rogers, P. H., Hawkins, A. D., Popper, A. N., Fay, R. R., Gray, M. D., 2016. Parvulescu revisited: Small tank acoustics for bioacousticians. Adv. Exp. Med. Biol. 875, 933–941. https://doi.org/10.1007/978-1-4939-2981-8_115/COV.

Ruiz-Ruiz, P. A., Hinojosa, I. A., Urzua, A., Urbina, M. A., 2020. Anthropogenic noise disrupts mating behavior and metabolic rate in a marine invertebrate. Proc. Meet. Acoust. 37(1), 040006. https://doi.org/10.1121/2.0001302.

Sal Moyano, M. P., Ceraulo, M., Hidalgo, F. J., Luppi, T., Nuñez, J., Radford, C. A., Mazzola, S., Gavio, M. A., Buscaino, G., 2021. Effect of biological and anthropogenic sound on the orientation behavior of four species of brachyuran crabs. Mar. Ecol. Prog. Ser. 669, 107–120. https://doi.org/10.3354/meps13739.

Sarà, G., Dean, J. M., D’Amato, D., Buscaino, G., Oliveri, A., Genovese, S., Ferro, S., Buffa, G., Lo Martire, M., Mazzola, S., 2007. Effect of boat noise on the behaviour of bluefin tuna *Thunnus thynnus* in the Mediterranean Sea. Mar. Ecol. Prog. Ser. 331, 243–253. https://doi.org/10.3354/MEPS331243.

Schaub, A., Ostwald, J., & Siemers, B. M., 2008. Foraging bats avoid noise. J. Exp. Biol. 211(19), 3174–3180. https://doi.org/10.1242/JEB.022863.

Scott, K. N., 2004. International Regulation Of Undersea Noise. Int. Comp. Law. Q. 53(2), 287–323. https://doi.org/10.1093/ICLQ/53.2.287.

Shafiei Sabet S., Neo, Y. Y., Slabbekoorn, H., 2015. The effect of temporal variation in sound exposure on swimming and foraging behaviour of captive zebrafish, Anim. Behav. 107, 49–60.

Shafiei Sabet, S., Karnagh, S. A., Azbari, F. Z., 2020. Experimental test of sound and light exposure on water flea swimming behaviour. Proc. Meet. Acoust. 38, 010015. https://doi.org/10.1121/2.0001270.

Shafiei Sabet, S., Van Dooren, D., Slabbekoorn, H., 2016a. Son et lumière: Sound and light effects on spatial distribution and swimming behavior in captive zebrafish. Environ. Poll. 212, 480–488. https://doi.org/10.1016/j.envpol.2016.02.046.

Shafiei Sabet, S., Wesdorp, K., Campbell, J., Snelderwaard, P., Slabbekoorn, H., 2016b. Behavioural responses to sound exposure in captivity by two fish species with different hearing ability. Anim. Behav. 116, 1–11. https://doi.org/10.1016/j.anbehav.2016.03.027.

Shannon, G., McKenna, M. F., Angeloni, L. M., Crooks, K. R., Fristrup, K. M., Brown, E., Warner, K. A., Nelson, M. D., White, C., Briggs, J., McFarland, S., Wittemyer, G., 2016. A synthesis of two decades of research documenting the effects of noise on wildlife. Biol. Rev. 91(4), 982–1005. https://doi.org/10.1111/BRV.12207.

Siemers, B. M., Schaub, A., 2011. Hunting at the highway: traffic noise reduces foraging efficiency in acoustic predators. Proc. Royal Soc. B: Biol. Sci. 278(1712), 1646–1652. https://doi.org/10.1098/RSPB.2010.2262.

Simpson, S. D., Radford, A. N., Holles, S., Ferarri, M. C. O., Chivers, D. P., McCormick, M. I., Meekan, M. G., 2016. Small-Boat Noise Impacts Natural Settlement Behavior of Coral Reef Fish Larvae. Adv. Exp. Med. Biol. 875, 1041–1048. https://doi.org/10.1007/978-1-4939-2981-8_129.

Simpson, S. D., Radford, A. N., Nedelec, S. L., Ferrari, M. C. O., Chivers, D. P., McCormick, M. I., Meekan, M. G., 2016. Anthropogenic noise increases fish mortality by predation. Nat. Commun. 7:1 7(1), 1–7. https://doi.org/10.1038/ncomms10544.

Slabbekoorn, H., Ripmeester, E. A. P., 2008. Birdsong and anthropogenic noise: implications and applications for conservation. Mol. Ecol. 17(1), 72–83. https://doi.org/10.1111/J.1365-294X.2007.03487.X.

Slabbekoorn, H., Bouton, N., van Opzeeland, I., Coers, A., ten Cate, C., Popper, A. N., 2010. A noisy spring: the impact of globally rising underwater sound levels on fish. Trends Ecol. Evol. 25(7), 419–427. https://doi.org/10.1016/J.TREE.2010.04.005.

Slabbekoorn, H., Dalen, J., de Haan, D., Winter, H. V., Radford, C., Ainslie, M. A., Heaney, K. D., van Kooten, T., Thomas, L., Harwood, J. 2019. Population-level consequences of seismic surveys on fishes: An interdisciplinary challenge. Fish Fish. 20(4), 653–685. https://doi.org/10.1111/FAF.12367.

Slater, M., Fricke, E., Weiss, M., Rebelein, A., Bögner, M., Preece, M., Radford, C., 2020. The impact of aquaculture soundscapes on whiteleg shrimp *Litopenaeus vannamei* and Atlantic salmon *Salmo salar*. Aquac. Environ. Interact. 12, 167–177.https://doi.org/10.3354/AEI00355.

Smith, M.E., Kane, A.S., Popper, A.N., 2004. Noise-induced stress response and hearing loss in goldfish (*Carassius auratus*). J. Exp. Biol. 207:427–435

Snitman, S. M., Mitton, F. M., Marina, P., Maria, C., Giuseppa, B., Gavio, M. A., Sal Moyano, M. P., 2022. Effect of biological and anthropogenic habitat sounds on oxidative stress biomarkers and behavior in a key crab species. Comp. Biochem. Physiol. C: Toxicology and Pharmacology. 257, 1–42. https://doi.org/10.1016/j.cbpc.2022.109344.

Solan, M., Hauton, C., Godbold, J. A., Wood, C. L., Leighton, T. G., White, P., 2016. Anthropogenic sources of underwater sound can modify how sediment-dwelling invertebrates mediate ecosystem properties. Sci. Rep. 6(1), 1–9. https://doi.org/10.1038/srep20540.

Southall, B. L., Bowles, A. E., Ellison, W. T., Finneran, J. J., Gentry, R. L., Greene, C. R., Kastak, D., Ketten, D. R., Miller, J. H., Nachtigall, P. E., Richardson, W. J., Thomas, J. A., Tyack, P. L., 2008. Marine mammal noise-exposure criteria: Initial scientific recommendations. Bioacoustics, 17(1-3), 273–275. https://doi.org/10.1080/09524622.2008.9753846.

Spanier, E., Tom, M., Pisants, S., Almong, G., 1988. Seasonality and Shelter Selection By The Slipper Lobate *Scyllarides latus* In The Southeastern Mediterranean. Mar. Ecol. Prog. Ser., 42:247–255.

Stocker, M., 2002. Fish, mollusks and other sea animals’ use of sound, and the impact of anthropogenic noise in the marine acoustic environment. J. Acoust. Soc. Am. 112(5), 24–31. https://doi.org/10.1121/1.4779979.

Tornero, V., Hanke, G., 2016. Chemical contaminants entering the marine environment from sea-based sources: A review with a focus on European seas. Mar. Pollut. Bull. 112(1-2), 17–38. https://doi.org/10.1016/J.MARPOLBUL.2016.06.091.

Ukaogo, P. O., Ewuzie, U., Onwuka, C. V., 2020. Environmental pollution: Causes, effects, and the remedies. Micro. Sust. Environ. Health, 419–429. https://doi.org/10.1016/B978-0-12-819001-2.00021-8.

Velilla, E., Collinson, E., Bellato, L., Berg, M. P., Halfwerk, W., 2021. Vibrational noise from wind energy-turbines negatively impacts earthworm abundance. Oikos, 130(6), 844–849. https://doi.org/10.1111/OIK.08166.

Voellmy, I. K., Purser, J., Flynn, D., Kennedy, P., Simpson, S. D., Radford, A. N., 2014. Acoustic noise reduces foraging success in two sympatric fish species via different mechanisms. Anim. Behav. 89(March), 191–198. https://doi.org/10.1016/j.anbehav.2013.12.029.

Wale, M. A., Simpson, S. D., Radford, A. N., 2013. Noise negatively affects foraging and antipredator behaviour in shore crabs. Anim. Behav. 86(1), 111–118. https://doi.org/10.1016/j.anbehav.2013.05.001.

Weber, S., Traunspurger, W., 2016. Influence of the ornamental red cherry shrimp *Neocaridina davidi*(Bouvier, 1904) on freshwater meiofaunal assemblages. Limnologica, 59(June 2016), 155–161. https://doi.org/10.1016/j.limno.2016.06.001.

Williams, R., Wright, A. J., Ashe, E., Blight, L. K., Bruintjes, R., Canessa, R., Clark, C. W., Cullis-Suzuki, S., Dakin, D. T., Erbe, C., Hammond, P. S., Merchant, N. D., O’Hara, P. D., Purser, J., Radford, A. N., Simpson, S. D., Thomas, L., Wale, M. A., 2015. Impacts of anthropogenic noise on marine life: Publication patterns, new discoveries, and future directions in research and management. Ocean Coast. Manag. 115, 17–24. https://doi.org/10.1016/JOCECOAMAN.2015.05.021.

Wowor, D., Cai, Y., Ng, P.K.L., 2004. Crustacean: Decapoda: Caridea. In: Yule, C., Yong, H.S. (Eds.), The Freshwater Invertebrates of Malaysia and Singapore. Malaysian Academy of Sciences, Kuala Lumpur, pp. 337–357.

